# Targeting Oncogenic Src Homology 2 Domain-Containing Phosphatase 2 (SHP2) by Inhibiting its Protein-Protein Interactions

**DOI:** 10.1101/2020.08.28.271809

**Authors:** Sara Bobone, Luca Pannone, Barbara Biondi, Maja Solman, Elisabetta Flex, Viviana Canale, Paolo Calligari, Chiara De Faveri, Tommaso Gandini, Andrea Quercioli, Giuseppe Torini, Martina Venditti, Antonella Lauri, Giulia Fasano, Jelmer Hoeksma, Valerio Santucci, Giada Cattani, Alessio Bocedi, Giovanna Carpentieri, Valentina Tirelli, Massimo Sanchez, Cristina Peggion, Fernando Formaggio, Jeroen den Hertog, Simone Martinelli, Gianfranco Bocchinfuso, Marco Tartaglia, Lorenzo Stella

## Abstract

We developed a new class of inhibitors of protein-protein interactions of the SHP2 phosphatase, which is pivotal in multiple signaling pathways and a central target in the therapy of cancer and rare diseases. Currently available SHP2 inhibitors target the catalytic site or an allosteric pocket but lack specificity or are ineffective on disease-associated SHP2 mutants. Based on the consideration that pathogenic lesions cause signaling hyperactivation due to increased SHP2 association with cognate proteins, we developed peptide-based molecules with low nM affinity for the N-terminal Src homology domain of SHP2, good selectivity, stability to degradation and an affinity for pathogenic variants of SHP2 up to 20 times higher than for the wild-type protein. The best peptide reverted the effects of a pathogenic variant (D61G) in zebrafish embryos. Our results provide a novel route for SHP2-targeted therapies and a tool to investigate the role of protein-protein interactions in the function of SHP2.

## Introduction

### SHP2 in physiology and pathology

Tyrosine phosphorylation, regulated by protein-tyrosine kinases (PTKs) and protein-tyrosine phosphatases (PTPs), is a fundamental mechanism of cell signaling. Aberrant tyrosine phosphorylation, caused by hyperactive PTKs, occurs in many malignancies and most current targeted anticancer drugs are PTK inhibitors. PTPs counteract the effects of kinases, and therefore they are generally considered negative regulators of cell signaling and tumor suppressors [Elson 2018]. However, the Src homology 2 (SH2) domain-containing phosphatase 2 (SHP2), encoded by the *PTPN11* gene, is a non-receptor PTP that does not conform to this simplistic picture [Tajan 2015].

SHP2 is ubiquitously expressed and mediates signal transduction downstream of various receptor tyrosine kinases (RTKs): it is required for full and sustained activation of the RAS/MAP kinase pathway [Saxton 1997] and modulates signaling also through the PI3K-AKT and JAK-STAT pathways, among others. Therefore, it is involved in the regulation of multiple cell processes, including proliferation, survival, differentiation, and migration [Tajan 2015]. Therefore, it is not surprising that dysregulated SHP2 function contributes to oncogenesis and underlies developmental disorders [Tajan 2015].

*PTPN11* was the first proto-oncogene encoding a tyrosine phosphatase to be identified [Tartaglia 2003]. Somatically acquired, gain of function mutations in *PTPN11* are the major cause of juvenile myelomonocytic leukemia (JMML), accounting for approximately 35% of cases [Tartaglia 2003]. JMML is a rare and aggressive myelodysplastic/myeloproliferative disorder of early childhood with a very poor prognosis, for which no drugs are presently available. Somatic *PTPN11* mutations also occur in childhood myelodysplastic syndromes, acute monocytic leukemia (AMoL, FAB M5) and acute lymphoblastic leukemia (ALL, “common” subtype) [Tartaglia 2003; 2004]. More rarely, activating mutations in this gene are found in adult myelodysplastic syndromes, chronic myelomonocytic leukemia, as well as solid tumors, including neuroblastoma, glioma, embryonal rhabdomyosarcoma, lung cancer, colon cancer and melanoma.

In addition to malignancies driven by *PTPN11* mutations, several forms of cancer are linked to the activity of wild type (WT) SHP2, too. By screening hundreds of cancer cell lines with a shRNA library, a landmark study showed that SHP2 is required for survival of RTK-driven cancer cells [Chen 2016]. SHP2 is also a central node in intrinsic and acquired resistance to targeted cancer drugs [Prahallad 2015], which is often caused by RTK activation through feedback loops.

SHP2 is a mediator of immune checkpoint pathways, such as PD-1 [Okazaki 2013]. These signaling cascades inhibit the activation of immune cells, thus allowing self-tolerance and modulation of the duration and amplitude of physiological immune responses. SHP2 binds to the activated receptors and is responsible for starting the signaling cascade that prevents immune cell activation [Okazaki 2013]. Some cancer cells are able to hijack these signaling pathways, thus evading antitumor immune defenses; therefore, SHP2 is currently being considered as a possible target for cancer immunotherapy [Marasco 2020a]. Finally, it is worth mentioning that induction of gastric carcinoma by *H. pylori* is mediated by the interaction of its virulence factor CagA with SHP2, causing aberrant activation of the phosphatase [Higashi 2002, Hayashi 2017].

In addition to its role in cancer, SHP2 is involved in a family of rare diseases collectively known as RASopathies. Germline missense mutations in *PTPN11* occur in ∼50% of individuals affected by Noonan syndrome (NS) [Tartaglia 2001], one of the most common non-chromosomal disorders affecting development and growth [Roberts 2013], and in ∼90% of patients affected by the clinically related Noonan syndrome with multiple lentigines (NSML, formerly known as LEOPARD syndrome) [Digilio 2002]. RASopathies are characterized by congenital cardiac anomalies, hypertrophic cardiomyopathy, short stature, musculoskeletal anomalies, facial dysmorphisms, variable intellectual disability and susceptibility to certain malignancies [Tartaglia 2010]. To date, the only treatment in use for NS and related disorders is growth hormone therapy, to improve linear growth [Roberts 2013].

### Structure and allosteric regulation of SHP2

The structure of SHP2 includes two Src homology 2 (SH2) domains, called N-SH2 and C-SH2, followed by the catalytic PTP domain, and an unstructured C-terminal tail (Figure 1) [Tajan 2015]. SH2 domains are recognition elements that bind protein sequences containing a phosphorylated tyrosine (pY) [Liu 2006, Anselmi 2020]. In SHP2, they mediate association to RTKs, cytokine receptors, cell adhesion molecules and scaffolding adaptors. Therefore, SHP2 (together with the closely related SHP1) is recruited (through its SH2 domains) by motifs containing two pYs and dephosphorylates other (or even the same) pYs through its PTP domain. The crystallographic structures of SHP2 [Hof 1998, LaRochelle 2018], complemented by biochemical analyses [Keilhack 2005; Tartaglia 2006; Bocchinfuso 2007; Martinelli 2008], have elucidated the main features of the allosteric regulation of SHP2 activity. Under basal conditions, the N-SH2 domain blocks the active site of the PTP domain, inserting a loop (DE or “blocking” loop) in the catalytic pocket. Consistently, the basal activity of SHP2 is very low. Association of SHP2 to its binding partners through the SH2 domains favors the release of this autoinhibitory interaction, making the catalytic site available to substrates, and causing activation (Figure 1). Specifically, structures of the N-SH2 domain associated to phosphopeptide sequences show that association to binding partners induces a conformational change in the blocking loop, which loses complementarity to the active site [Lee 1994]. At the same time, the N-SH2/PTP interaction allosterically controls the conformation of the N-SH2 domain binding site. Structures of the autoinhibited protein show that the binding site of the N-SH2 domain is closed by two loops (EF and BG). By contrast, in structures of the isolated N-SH2 domain [Lee 1994], or the recently reported structure of the active state of SHP2 [LaRochelle 2018], the binding site is open (Figure 1). Consequently, we and others have hypothesized that the transition between the closed, autoinhibited state and the open, active conformation is coupled to an increased affinity for binding partners [Keilhack 2005; Bocchinfuso 2007, Martinelli 2008, LaRochelle 2018].

**Figure 1:**
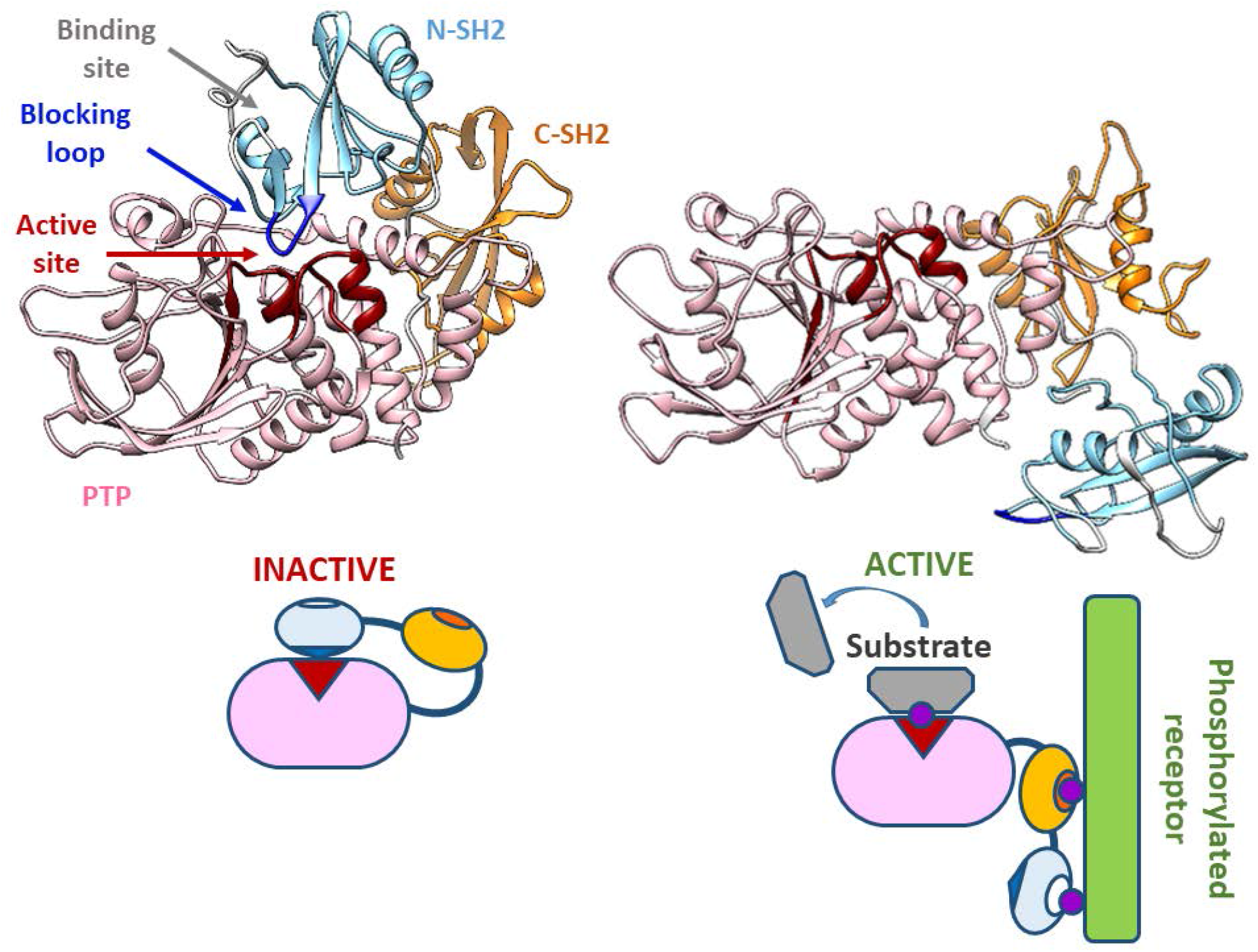
SHP2 structure and scheme of the activation process. **Top**: crystallographic structures for the closed, auto-inhibited and the open, active states of SHP2 (left and right, respectively). The N-SH2, C-SH2 and PTP domain are colored in light blue, orange and pink, respectively. The N-SH2 blocking loop (DE loop) is colored in blue, while the PTP active site is highlighted in dark red. The EF and BG N-SH2 loops, controlling access to the binding site for phosphorylated sequences of binding partners, are represented in white. PDB codes of the two structures are 2SHP and 6CRF. Segments missing in the experimental structures were modeled as previously described [Bocchinfuso 2007]. **Bottom**: schematic model of the allosteric regulation mechanism.

The spectrum of pathogenic *PTPN11* mutations is generally consistent with this picture of SHP2 regulation. Most mutations cluster at the N-SH2/PTP interface, destabilizing the interaction between these two domains and causing constitutive activation of the phosphatase [Keilhack 2005; Tartaglia 2006; Bocchinfuso 2007]. These mutations concomitantly induce an increased responsiveness to activation by association of bisphosphorylated sequences to the SH2 domains [Keilhack 2005; Bocchinfuso 2007; Martinelli 2008; LaRochelle 2018]. Other mutations localize in the binding site of the SH2 domains, and simply cause an increased affinity for phosphorylated binding partners [Tartaglia 2006]. In all cases, the final effect is an upregulation of the RAS/MAPK signal transduction pathway.

### SHP2 as a pharmacological target

All the findings reported above clearly indicate SHP2 as an important molecular target for cancer and RASopathies [Tang 2020]. Since SHP2 is a convergent node for multiple signaling pathways, SHP2 inhibitors may represent a way to suppress the effect of disease-causing mutations involving different proteins along the signaling cascade with a single molecule [Mullard 2018].

Research efforts in SHP2-targeted drug discovery have long been focused mainly on active-site inhibitors [Yuan 2020, Mostinski 2020]. Several molecules inhibiting the catalytic activity of SHP2 have been reported, but many of them are affected by the same limitations that led PTPs in general to be considered “undruggable”, *i.e.* lack of target specificity and poor bioavailability [Mullard 2018, Yuan 2020]. Some compounds with good affinity and apparent selectivity have been described, but more recent studies demonstrated that these molecules have several off-target effects [Tsutsumi 2018].

An alternative pharmacological strategy has been pursued by researchers at Novartis [Chen 2016, Garcia Fortanet 2016, Bagdanoff 2019, Sarver 2019], followed by others [Xie 2017, Mullard 2018; Wu 2018; Tang 2020], who reported allosteric inhibitors stabilizing the autoinhibited structure of SHP2 by binding to a pocket located at the interdomain interface in the closed conformation of the phosphatase. SHP099, the inhibitor developed by Novartis, is finding promising applications in the treatment of RTK-driven cancers [LaMarche 2020] and in combined therapy against drug resistant cells [Prahallad 2015]. These results have spurred a renewed interest in the inhibition of phosphatases [Mullard 2018]. Currently, several allosteric inhibitors stabilizing the closed conformation of SHP2 are undergoing clinical trials [Tang 2020]: TNO155 [LaMarche 2020] by Novartis (derived from SHP099), RMC-4630, by Revolution Medicines and Sanofi, JAB-3068 and JAB-3312, developed by Jacobio Pharmaceuticals and RLY-1971, by Relay Therapeutics. Allosteric inhibitors have also been used to target SHP2 to proteolytic degradation [Wang 2020]. However, these compounds are generally poorly effective in the case of activating *PTPN11* mutants, since the allosteric binding site is lost in the open conformation of the enzyme [LaRochelle 2018, Tang 2020].

### Inhibition of protein-protein interactions as an alternative pharmacological strategy

Due to the allosteric mechanism described above, SHP2 activation and its association to binding partners are coupled events. Therefore, the effect of NS- and leukemia-causing mutations destabilizing the autoinhibited conformation is twofold: they cause an increase in the phosphatase activity of the protein, but at the same time favor the N-SH2 conformation suitable for binding phosphorylated proteins, thus increasing the overall responsiveness of SHP2 to its interaction partners. Several lines of evidence indicate that the second event, rather than the enhanced basal activity, is essential for the abnormal activation of the RAS/MAPK pathway.

Some pathogenic mutations, such as the NS-associated p.T42A, simply increase the binding affinity of the N-SH2 domain, without causing basal activation [Martinelli 2008, Keilhack 2005]; on the other hand, the ability of SHP2 to associate to binding partners is preserved in all the disease- associated *PTPN11* mutations [Tartaglia 2006, Martinelli 2012, 2020].

Truncated constructs with deletion or partial deletion of the N-SH2 domain cause a dramatic increase in the enzymatic activity of SHP2 and, at the same time, a complete loss of its ability to bind signaling partners. These constructs do not affect development in heterozygous mice [Saxton 1997] and do not cause any aberrant phenotype in cells [Saxton 1997, Higashi 2002]. Indeed, RAS/MAPK signaling in homozygous cells with the mutated construct was reduced with respect to the WT cells [Shi 1998]. However, cellular morphological changes (hummingbird phenotype) were observed when the truncated construct was targeted to cellular membranes by adding a membrane-localization signal [Higashi 2002], demonstrating the importance of proper cellular localization, normally mediated by SH2 domains. The relevance of SHP2 association to its binding partners for its role in aberrant signaling has been demonstrated also by a study on monobodies targeting the N-SH2 domain and disrupting its association with adaptor proteins. Expression of these monobodies in cancer cells carrying the activating *p.*V45L mutation abolished ERK1/2 phosphorylation almost completely [Sha 2013]. Similarly, Kertész and coworkers [Kertész 2006] reported that the natural SHP2-binding motif of Gab1, when delivered into immune cells, modulated phosphorylation patterns.

An example of the opposite situation, where binding is preserved and the catalytic activity is impaired, is provided by *PTPN11* mutations causing NSML, such as T468M. This class of amino acid substitutions are located in proximity of the PTP active site, at the PTP/N-SH2 interface, and have a twofold effect: they destabilize the closed state of the protein, and consequently promote SHP2 association to signaling partners; at the same time, they perturb the active site and therefore strongly impair the catalytic activity of the phosphatase. Interestingly, the phenotype of NSML is very similar to that of NS, and these mutations still allow the activation of multiple effector pathways [Martinelli 2008; Yu 2014].

Overall, these findings strongly suggest that a mere enhancement in SHP2 catalytic activity is not sufficient to cause disease and indicate that increased association to binding partners plays a major role in the pathogenic mechanism associated with *PTPN11* mutations. Therefore, inhibition of SHP2 binding to other proteins through its SH2 domains represents a promising alternative pharmaceutical strategy. No molecules targeting the SH2 domains of SHP2 for therapeutic purposes have been developed so far, even though SH2 domains in general have received much attention as potential pharmaceutical targets [Machida 2005, Cerulli 2020]. These recognition units generally have only moderate affinity and selectivity for cognate phosphorylated sequences, with dissociation constants in the 0.1 – 10 μM range [Kuriyan 1997, Machida 2005, Wagner 2013, Marasco 2020b]. However, we recently characterized the structural determinants of phosphopeptide binding by the N-SH2 domain of SHP2 [Anselmi 2020], and our data indicate this particular domain as a favorable exception, since its peculiar features make significantly higher affinities possible.

Based on these considerations, we explored the possibility to target SHP2 protein-protein interactions (PPIs), rather than its catalytic activity. We developed a peptide-based molecule with low nM affinity to the N-SH2 domain of SHP2, high specificity and resistance to degradation. This inhibitor rescued the mortality and developmental defects induced by a pathogenic mutation *in vivo*. Our results provide a novel route for SHP2-targeted therapies and offer a new tool to further investigate the role of SHP2 PPIs in the signaling cascades controlled by this phosphatase.

## Results

### 1) Characterization of IRS-1 pY1172/N-SH2 binding

#### 1.1) The IRS-1 pY1172 peptide binds the N-SH2 domain with a low nanomolar affinity

The peptide corresponding to pY 1172 (rat sequence, SLN-pY-IDLDLVKD) or pY 1179 (human sequence, GLN-pY-IDLDLVKD) of insulin receptor substrate 1 (IRS-1) has one of the highest known binding affinities for the N-SH2 domain of SHP2 [Anselmi 2020, Marasco 2021]. Based on our study of the structural determinants of high binding affinity to this domain, the IRS-1 pY 1772 sequence is near to optimal under several respects, since it has apolar residues at positions +1, +3 and +5, which point towards the hydrophobic groove in the N-SH2 structure, and anionic amino acids at positions +2 and +4, which can interact with a peculiar KxK motif in the BG loop [Anselmi 2020].

The binding affinity of the IRS-1 pY1172 peptide has been characterized in several literature studies. Unfortunately, these results are extremely contradictory, as reported in Table S1, with dissociation constants ranging from ∼10 nM to ∼10 µM. Several possible factors can be invoked to explain these discrepancies, including the effect of radioactive labels [Case 1994], dimerization of GST-N-SH2 constructs [Sugimoto 1994, Ladbury 1995] even at low nM concentrations [Fabrini 2009], or the sensitivity of the technique [Kelihack 2005]

**Table 1.** Peptide sequences investigated in this study.

| Abbreviation | Sequence |
| --- | --- |
| P9 | GLN-pY-IDLDL |
| P9Y0 | GLN- Y-IDLDL |
| P8 | LN-pY-IDLDL |
| P7 | N-pY-IDLDL |
| P8W5 | LN-pY-IDLDW |
| P8F5 | LN-pY-IDLDF |
| P8E4W5 | LN-pY-IDLEW |
| P9ND0W5<br>(or OP) | GLN-ND-IDLDW |
| CF-P9 | CF-GLN-pY-IDLDL |
| CF-P9Y0 | CF-GLN- Y-IDLDL |
| CF-P9W5 | CF-GLN-pY-IDLDW |
| Cy3-P9W5 | Cy3-GLN-pY-IDLDW |
| CF-P9E4W5 | CF-GLN-pY-IDLEW |
| CF-P9ND0W5<br>(or CF-OP) | CF-GLN-ND-IDLDW |
| P8W5-TAT | LN-pY-IDLDW-GRKKRRQRRR |
| CF-P9W5-TAT | CF-GLN-pY-IDLDW-GRKKRRQRRR |

Considering these difficulties, in the present study, we developed a fluorescence anisotropy binding assay. In a direct binding experiment, the fluorescently labeled peptide IRS-1 pY1172 analog CF- P9 (Table 1) was titrated with increasing concentrations of the N-SH2 domain. The fraction of protein-bound peptide was determined from the increase in fluorescence anisotropy (Figure 2), and a *Kd* of 53±2 nM was obtained (Table 2).

**Figure 2:**
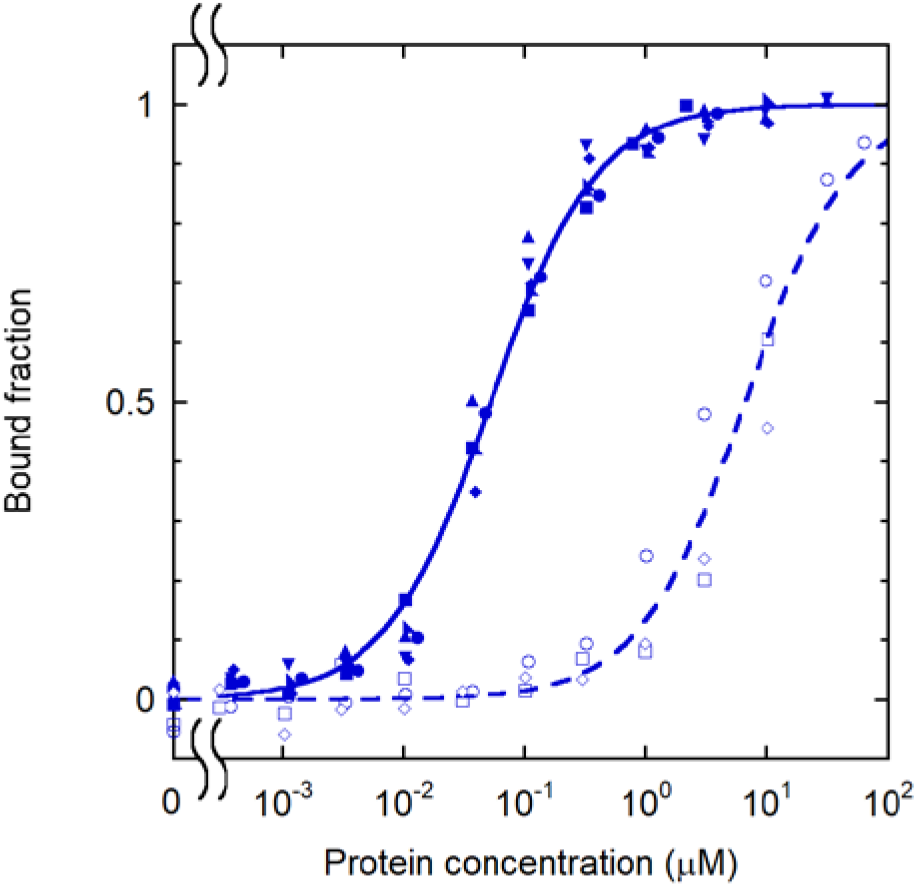
Binding curves for the phosphorylated and unphosphorylated IRS-1 pY1172 peptides. [CF-P9]=1.0 nM (full symbols and solid line), [CF-P9Y0]=10 nM (empty symbols and dashed line); Replicate experiments are reported with different symbols and were fit collectively.

**Table 2.** Dissociation constants obtained from the fluorescence anisotropy binding experiments.

| Peptide | Domain/Protein | $K_d$ (nM) |
| --- | --- | --- |
| CF-P9 | N-SH2 | 53 ± 2 |
| CF-P9Y0 | N-SH2 | 6600 ± 600 |
| CF-P9W5 | N-SH2 | 3.3 ± 0.2 |
| CF-P9W5 | C-SH2 | 4200 ± 300 |
| Cy3-P9W5 | N-SH2 | 23 ± 2 |
| CF-P9E4W5 | N-SH2 | 8.2 ± 0.7 |
| CF-P9ND0W5 (CF-OP) | N-SH2 | 68 ± 5 |
| CF-P9ND0W5 (CF-OP) | PTP | 10000 ± 800 |
| CF-P9ND0W5 (CF-OP) | SHP2 (WT) | 930 ± 70 |
| CF-P9ND0W5 (CF-OP) | SHP2 (A72S) | 400 ± 40 |
| CF-P9ND0W5 (CF-OP) | SHP2 (E76V) | 330 ± 10 |
| CF-P9ND0W5 (CF-OP) | SHP2 (D61H) | 170 ± 10 |
| CF-P9ND0W5 (CF-OP) | SHP2 (F71L) | 140 ± 10 |
| CF-P9ND0W5 (CF-OP) | SHP2 (E76K) | 48 ± 2 |

All peptides were amidated at the C-terminus. Unlabeled peptides were acetylated at the N- terminus. CF is 5,6 carboxyfluorescein, Cy3 is Cyanine 3 carboxylic acid and ND is the non- dephosphorylatable pY mimic phosphonodifluoromethyl phenylalanine (F_2_Pmp). The optimized peptides are highlighted in grey. RP-HPLC retention times (R_t_), purities, theoretical molecular weights and those determined experimentally by ESI-MS spectrometry are reported in Table S2.

#### 1.2) Phosphorylation contributes only 30% of the standard binding free energy

Association of SH2 domains with the partner proteins is regulated by phosphorylation, and therefore the phosphate group is necessarily responsible for a large fraction of the binding affinity. On the other hand, to have a good selectivity, the rest of the sequence must also contribute significantly to the peptide/protein interaction. To quantify this aspect, we performed a binding experiment (Figure 2) with an unphosphorylated analog of the labeled IRS-1 pY1172 peptide, CF- P9Y0 (Table 1). The affinity was approximately 100 times lower, with a *K_d_* of 6.6±0.6 μM, compared with 53±2 nM for the phosphorylated peptide. The corresponding values for the standard free energy of binding (assuming a 1 M standard state) are -29.6±0.2 kJ/mol and -41.6±0.1 kJ/mol, respectively. Assuming additivity of contributions, the phosphate group results to be responsible for the difference of -12.0±0.2 kJ/mol, *i.e.* for less than 30% of the total standard binding free energy of the phosphorylated peptide. This result indicates that the contribution of the rest of the peptide predominates in the binding interactions and bodes well for our design efforts.

### 2) Sequence optimization

#### 2.1) The sequence can be reduced to 8 amino acids without loss in affinity

Literature data are partially contradictory regarding the effect of shortening the IRS-1 pY1172 sequence on the binding affinity. Kay [1998] reported that the sequence could be shortened at the C-terminus down to the +5 residue and at the N-terminus down to the -2 position, without any loss in affinity. By contrast, Case [1994] observed a significant reduction in affinity by shortening the sequence from SLN-pY-IDLDLVKD to LN-pY-IDLDLV. Our previous study clearly indicated that residues -2 to +5 are the most important for the interaction [Anselmi 2020]. To clarify the role of N-terminal residues in determining the N-SH2 domain binding affinity, we performed displacement studies (Figure S1) with the unlabeled peptide P9, and with the shortened analogues P8 and P7 (Table 1), where residues -3 or -2 and -3 were removed, respectively. No significant loss in affinity was observed by reducing the sequence to 8 residues, while removal of the amino acid at position -2 caused a drastic perturbation of complex stability (Figure S1). The -2 to +5 IRS-1 sequence is the minimal peptide with a low nM dissociation constant.

#### 2.2) Single amino acids substitutions improve the *K_d_* to the low nM range

Hydrophobic residues are required at position +1, +3 and +5 of the phosphopeptide sequence [Anselmi 2020], but aromatic residues are present in some natural high affinity binding sequences, at position +5 only [Case 1994, Huyer 1995; Hayashi 2017, Bonetti 2018, Marasco 2020a]. The crystallographic structures of some of these complexes [Lee 1994, Hayashi 2017, Marasco 2020] show that an aromatic side chain can be accommodated by a relatively large hydrophobic pocket and that the +5 peptide residue interacts with the BG and EF loops of the domain, which are important for binding specificity [Lee 1994, Anselmi 2020]. Finally, a preference for aromatic residues at position +5 has been indicated by several peptide library studies [De Souza 2002; Imhof 2006; Martinelli 2012; Tinti 2013].

Based on these considerations, we analyzed in silico the effect of different aromatic amino acids at position +5. Free energy calculations indicated that substitution of L with the bulkier W (but not with F) could be favorable (Figure 3). The additional substitution of D in +4 with the longer E was evaluated as well, in view of a possible strengthening of the electrostatic interactions with the KxK motif in the BG loop. However, in this case no further increase in binding affinity was predicted by the free energy calculations (Figure 3).

**Figure 3:**
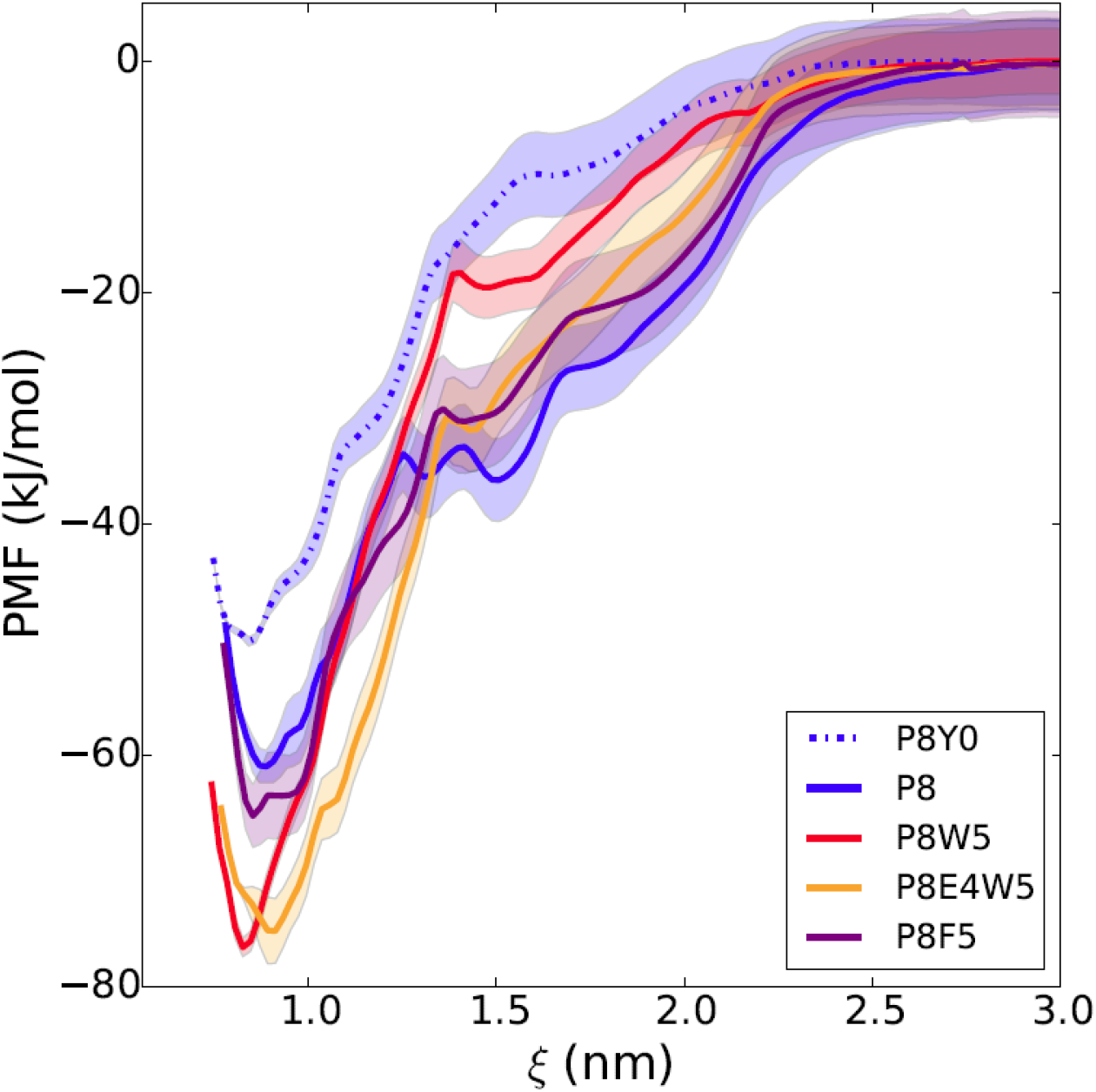
in silico free energy calculations for different modified sequences. The free energy profile is reported as a function of the distance between the centers of mass of the N-SH2 domain and of the phosphopeptide. The simulations predict a loss in affinity of P8 (blue line) with dephosphorylation of the pY (dashed blue line), and a gain with substitution of the L in position +5 with W (red line), but not with F (violet line). The additional substitution of D in +4 with E (orange) does not provide any further increase in affinity. Shaded areas correspond to standard deviations in the PMF profile.

Analogs with F or W at position +5 (P8F5 and P8W5), as well as a labeled analog with the L to W substitution (CF-P9W5, Table 1) were synthesized and studied experimentally (Figure 4). As predicted, introduction of W in +5 was highly favorable, leading to reduction in the dissociation constant by an order of magnitude, both for the labeled and unlabeled analog (Tables 2 and 3). Consequently, the dissociation constant for the P8W5 analog was 1.5±0.3 nM. By contrast, the additional D to E substitution resulted in a slight loss in binding affinity (Figure 4 and Table 2 and 3).

**Figure 4:**
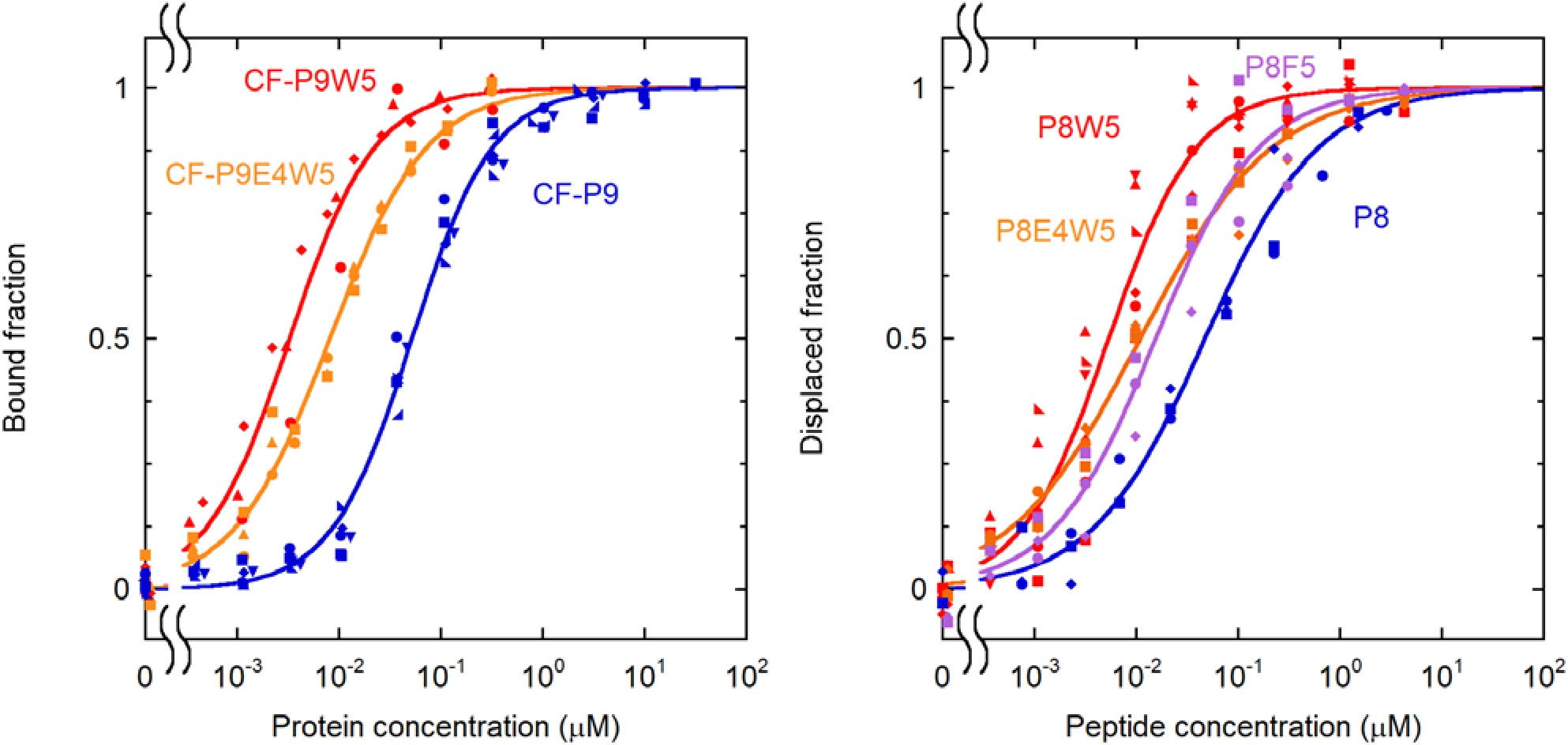
effect of substitutions at position +5 on the binding affinity. **Left**: direct binding experiments; [CF-P9W5]=0.10 nM, [CF-P9E4W5]=0.10 nM, [CF-P9]=1.0 nM. Data for CF-P9 are repeated here for comparison. **Right**: displacement assay, [CF-P9W5]=0.10 nM, [N-SH2]=3.35 nM.

Based on these results, further studies concentrated on the peptide with W at position +5.

### 3) Binding selectivity

#### 3.1) The modified sequence is highly selective for the N-SH2 domain of SHP2

The selectivity of binding of CF-P9W5 was first assessed with respect to the C-SH2 domain of SHP2, again with the fluorescence anisotropy assay (Figure S2). As reported in Table 2, the affinity for the C-SH2 domain was almost 1,000 times less than that for the N-SH2 domain.

A more complete analysis of the binding selectivity was performed on a protein array of 97 human SH2 domains (Figure 5). An analogue of CF-P9W5 was employed in this assay, where CF was substituted with the Cy3 dye, suitable for detection in the array reader. Control binding experiments showed that the change in fluorophore did affect peptide binding affinity only marginally (Table 2). Strikingly, significant binding was observed only with the N-SH2 domain of SHP2, and, to a lesser extent, to the SH2 domain of the adapter protein APS (also called SHP2B2). It is worth noting that binding to the N-SH2 domain of SHP1, which has the highest identity with that of SHP2 [Liu 2006], was negligible.

**Figure 5:**
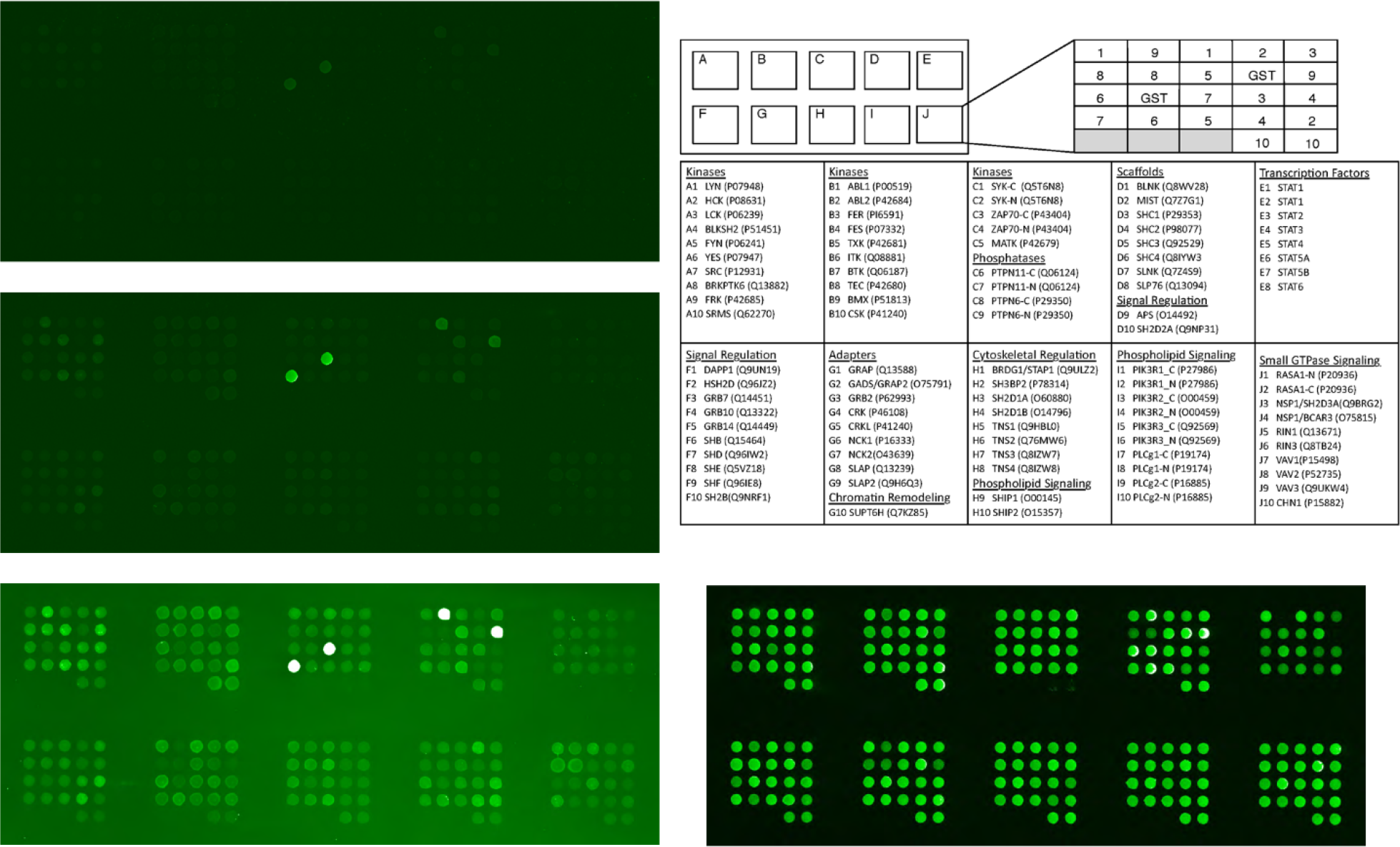
binding selectivity of Cy3-P9W5 for an array of SH2 domains. **Left**: fluorescence of the bound peptide, at a concentration of 0.5 nM (top), 5.0 nM (center) and 50 nM (bottom). Each SH2 domain was spotted in duplicate, and negative control spots (with GST only) are also present. The bright spots correspond to the N-SH2 domain of SHP2 and to the SH2 domain of the SH2 and PH domain-containing adapter protein APS (also called SHP2B2). The intensity of all other spots is comparable to that of the negative controls. **Right**: position of each SH2 domain in the array (top), and control of the protein loading in each spot, performed with an anti-GST antibody (bottom).

### 4) Engineering resistance to degradation

#### 4.1) Introduction of a non-hydrolysable pY mimic is compatible with low nM binding affinity

In view of intracellular or *in vivo* applications of the phosphopeptide, it is essential to make it resistant to degradation. The most labile moiety is the phosphate group of the pY residue, which can be hydrolyzed by protein tyrosine phosphatase, possibly also including SHP2, of which IRS-1 pY 1172 has been shown to be a substrate [Noguchi 1994]. We substituted the pY with the non- hydrolysable mimetic phosphonodifluoromethyl phenylalanine (F_2_Pmp), which is isosteric with pY and has a total negative charge comparable to that of pY under physiologic pH conditions [Burke 2006, Cerulli 2020].

Binding experiments demonstrated that the substituted analogue (CF-P9ND0W5, where ND0 indicates the introduction of the non-dephosphorylatable pY mimic at position 0, Table 1) has a dissociation constant for the N-SH2 domain that is just an order of magnitude worse with respect to that of CF-P9W5 (68 ± 5 nM with respect to 4.6 ± 0.4 nM) (Figure S3 and Table 2). Similarly, the dissociation constant for the unlabeled peptide P9ND0W5 was 15 ± 0.4 nM (with respect to 1.6 ± 0.4 nM for P8W5) and thus remained in the nM range (Table 3).

**Table 3.** Dissociation constants obtained from the displacement experiments. All measurements were performed on the N-SH2 domain of SHP2. Experiments were performed at [N-SH2]= 3.4 nM and [CF-P9W5]=0.5 nM (for P8 and P9ND0W5) or 0.1 nM (for the other peptides).

| Peptide | IC<sub>50</sub> (nM) | $K_i$ (nM) |
| --- | --- | --- |
| P8 | 47 ± 4 | 22 ± 4 |
| P8F5 | 16 ± 1 | 7 ± 2 |
| P8W5 | 5.4 ± 0.3 | 1.5 ± 0.3 |
| P8E4W5 | 11 ± 1 | 4 ± 2 |
| P9ND0W5 (OP) | 32 ± 5 | 14 ± 5 |

For the sake of brevity, in the following text, CF-P9ND0W5 and its unlabeled analogue P9ND0W5 will be also referred to as the optimized peptides, or CF-OP and OP, respectively.

#### 4.2) The optimized peptide OP is resistant to proteolytic degradation

To test the resistance to proteases, the optimized peptide OP was incubated in human serum for up to 90 minutes, or in DMEM for three days, and then analyzed by HPLC. No significant degradation was observed in these time frames (Figure S4). By contrast, the octadecapeptide HPA3NT3 [Park 2008], which we used as a positive control, was completely degraded already after 5 minutes (data not shown). This result bodes well for potential in vivo applications of the peptide.

### 5) Binding to and activation of the SHP2 protein

#### 5.1) OP binds to pathogenic mutants with much higher affinity than to the WT protein

As discussed in the introduction, we and others have hypothesized that, in its autoinhibited state, the conformation of the N-SH2 domain prevents efficient association to binding partners, while SHP2 binding affinity to phosphorylated sequences is maximized in the open, active state [Keilhack 2005; Bocchinfuso 2007, Martinelli 2008, LaRochelle 2018]. This model has many relevant consequences, because it implies that pathogenic mutants have a twofold effect: they increase the activity of the phosphatase, but also its affinity towards binding partners. In principle, both effects could be the origin of the hyperactivation of the signal transduction pathways involved in the pathologies caused by *PTPN11* mutations.

Notwithstanding the relevance of this aspect, to the best of our knowledge, no direct phosphopeptide binding experiments to the whole SHP2 protein have ever been performed, possibly due to the fact that pY can be dephosphorylated by the PTP domain. Now, OP and its fluorescent analogue CF-OP allow us to directly assess the hypothesis described above. Figure 6 and Table 2 report the results of binding experiments performed with CF-OP and WT SHP2 or the pathogenic mutants A72S, E76K, D61H, F71L and E76V. E76K is among the most common somatic lesions associated with leukemia and has never been observed as germline event in individuals with NS [Tartaglia 2003, 2006], as it results in early embryonic lethality [Xu 2011]. This mutation is strongly activating, with a basal activity of the corresponding mutant being at least 10 times higher than that of the WT protein. Conversely, A72S is a germline mutation specifically recurring among subjects with NS. In this case, basal activation is only twofold [Bocchinfuso 2007]. The D61H, F71L and E76V amino acid substitutions have been identified as somatic events in JMML and other leukemias [Tartaglia 2003] and, when transmitted in the germline, they are associated with a high prenatal lethality (M. Zenker, personal communication, 9/2019). Interestingly, we observed that the affinity for CF-OP nicely parallels the basal activity of these mutants (Figure 6). This finding provides a first direct confirmation that the closed, autoinhibited state has a lower affinity for the binding partners, compared to the open, active conformation.

**Figure 6:**
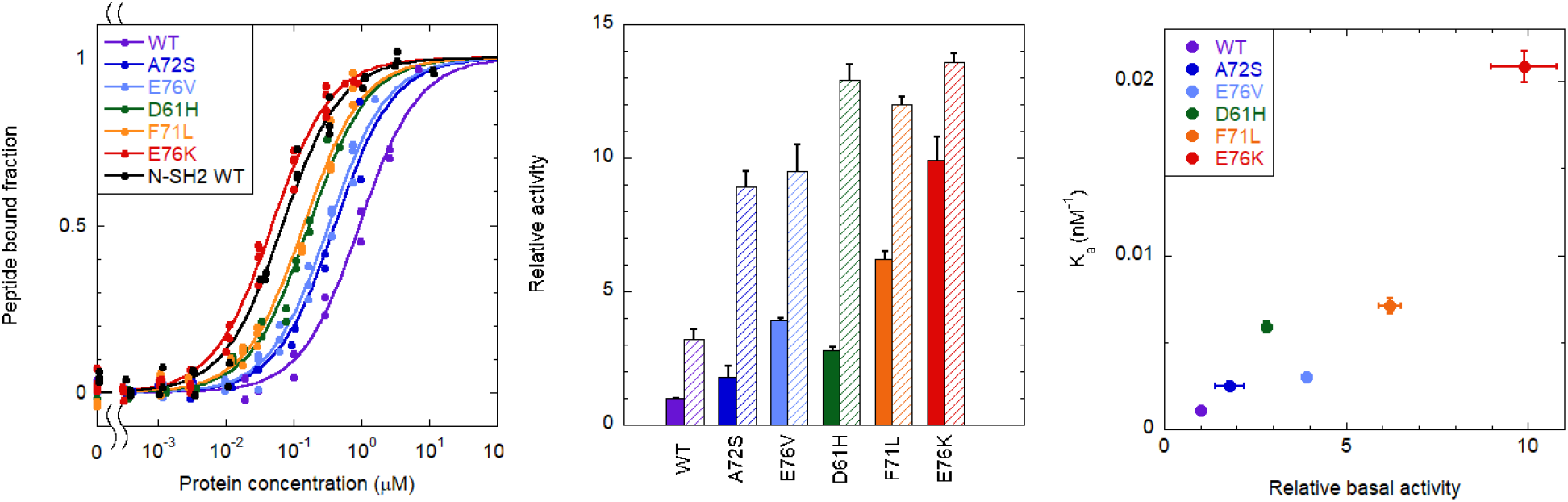
Binding of the CF-OP peptide to the whole SHP2 protein (WT and pathogenic mutants) **Left**: fraction of CF-OP peptide bound to the WT protein and selected mutants, obtained from fluorescence anisotropy experiments. The peptide bound fractions were obtained by following the variation of the peptide fluorescence anisotropy during the titration with increasing amounts of protein. [CF-OP]=1.0 nM. **Center**: catalytic activity of the WT protein and selected mutants, under basal conditions (solid bars) and after stimulation with 10 μM BTAM peptide (dashed bars). N=3 **Right**: correlation between basal activity and affinity (association constant). Error bars represent standard deviations.

#### 5.2) OP is also an inhibitor of the PTP domain

Based on previous reports of the dephosphorylation of IRS-1 pY 1172 by SHP2 [Noguchi 1994], we verified if P8 and P8W5 are also a substrate of this protein. These experiments were performed with a truncated SHP2 construct lacking the N-SH2 domain (SHP2_Δ104_), as it is fully activated and was shown to be more stable and less prone to aggregation compared to the isolated PTP domain [Martinelli 2020]. As reported in Figure S5, dephosphorylation was indeed observed, although to a lesser extent than for other phosphopeptides.

Using the non-dephosphorylatable peptide CF-OP, we measured directly binding to the PTP domain of SHP2 (Figure S6 and Table 2). Significant association was observed, although with a much lower affinity than with the N-SH2 domain (*K_d_* = 10.0 ± 0.8 μM). This finding indicates that in principle, the non-dephosphorylatable OP could act as a double hit SHP2 inhibitor, acting on both PPIs and catalytic activity.

#### 5.3) OP activates SHP2 only weakly

Binding of mono- or bi-phosphorylated peptides causes activation of SHP2. We tested the effect of OP on the WT protein, or on the A72S mutant (this experiment is not possible with E76K, as in that case the protein is essentially fully activated also in the absence of phosphopeptides). As shown in Figure S7, activation was very weak, compared with that induced by the bisphosphorylated BTAM peptide, and a significant effect was observed only with the mutant protein. Interestingly, under the experimental conditions used (10 μM peptide), the N-SH2 domain of both the WT and the A72S proteins is nearly saturated by the OP, according to the binding experiments reported in Figure 6. This finding indicates that the peptide could inhibit SHP2 PPIs, causing only a limited increase in catalytic activity. In any case, as demonstrated by studies on truncated constructs lacking the N-SH2 domain [Saxton 1997, Shi 1998, Higashi 2002], activation of SHP2 without proper PPIs has no pathogenic effects.

### 6) OP effectively reverses the effects of D61G mutation in vivo

We used the zebrafish model system to explore the in vivo effect of the peptide. Zebrafish Shp2a is highly homologous to human SHP2 (91.2% protein sequence identity); in particular, the sequence of the N-SH2 domain and of the N-SH2/PTP interface are identical in the human and fish proteins. RASopathies-associated mutants, including activating mutants of Shp2a, greatly impact zebrafish development. In humans, the D61G substitution has been found in both NS and leukemia [Kratz 2005] and in animal models it induces both NS-like features and myeloproliferative disease [Araki 2004]. Microinjection of synthetic mRNA encoding NS-associated mutants of Shp2 at the one-cell stage induces NS-like traits [Jopling 2007]. During gastrulation, convergence and extension movement are affected, resulting in oval-shaped embryos, with increased major/minor axis length ratio at 11 hpf [Jopling 2007]. Here we co-injected Shp2a-D61G mRNA with OP in zebrafish embryos, to investigate whether OP rescues the defective cell movements during gastrulation.

As shown in Figure 7 and Figure S8, we observed a dose-dependent decrease in Shp2a-D61G- induced major/minor axis ratios, with a rescue of the phenotype that was significant at 5 µM peptide concentration. On the other hand, embryos injected with Shp2a-WT were almost perfect spheres at 11 hpf, and co-injection with 3 µM peptide had no impact on their shape. As expected, a large portion of Shp2a-D61G injected embryos were severely affected and they died during embryonic development, whereas injection of WT Shp2a did not induce significant lethality. We followed the survival of Shp2a-D61G injected embryos and observed a significant and dose dependent improvement in the survival of embryos upon co-injection with 0.3 µM, 3 µM and 5 µM OP (Figure 8). By contrast, lethality of WT Shp2a embryos was not affected by co-injection of 3 µM OP.

**Figure 7:**
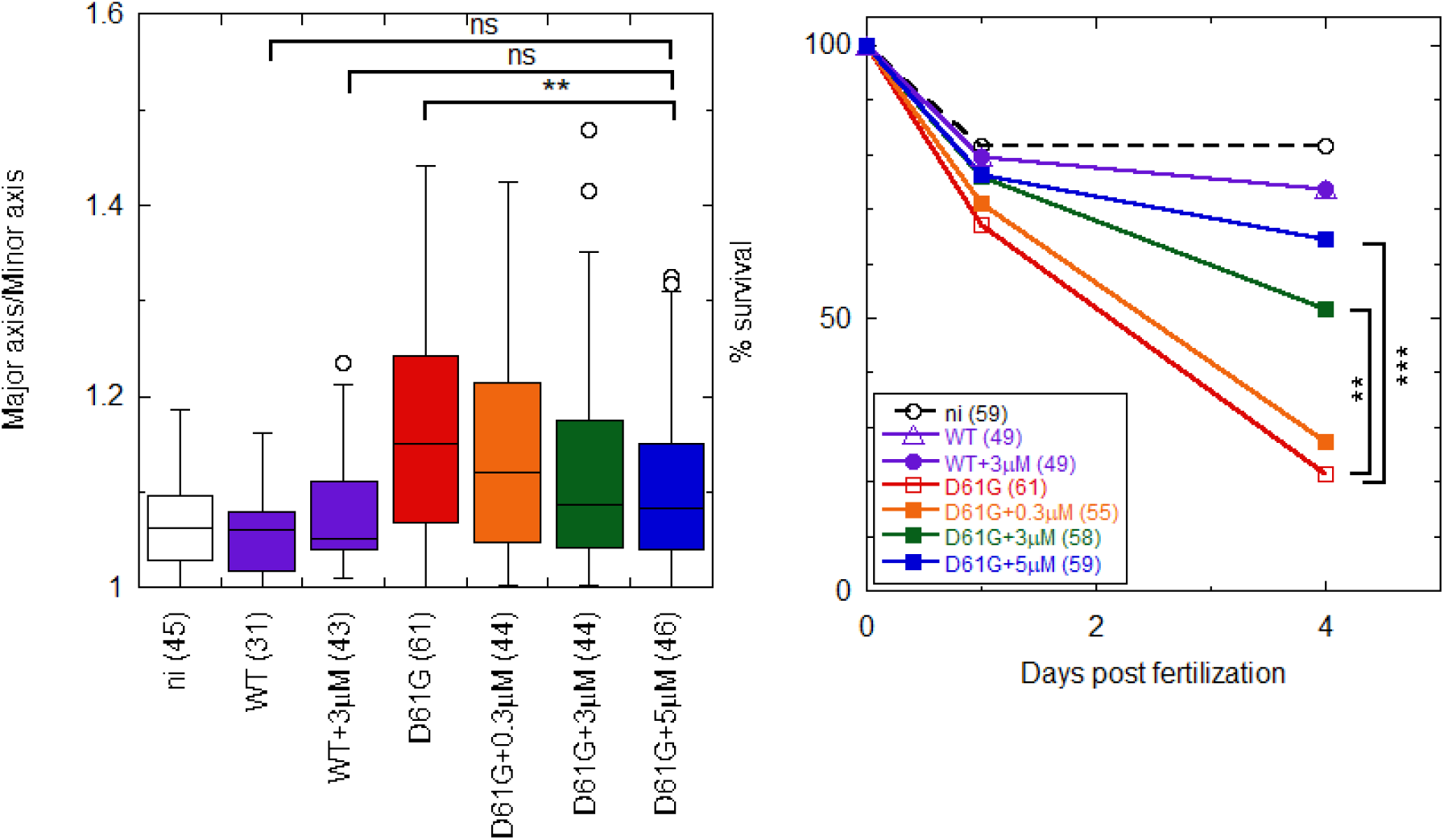
the OP partially rescues Shp2a-D61G-induced gastrulation defects and mortality in a dose dependent manner in zebrafish embryos. Embryos were injected at the one-cell stage with mRNA encoding GFP-2A-Shp2-D61G or GFP- Shp2-wt with or without peptide at 0.3 µM, 3 µM and 5 µM concentration. Non-injected embryos (ni) were evaluated as a control. **Left**: ovality of embryos at 11 hpf, as indicated by the ratio of the long and the short axis. Tukey’s honest significant difference test was done to assess significance. In the box plot, the horizontal line indicates the median, box limits indicate the 25^th^ and 75^th^ percentiles (interquartile range), whiskers (error bars) extend to the maximum and minimum values, or to 1.5 times the interquartile range from the 25^th^ and 75^th^ percentiles, if some data points fall outside this range. In the latter case, outliers are indicated as single data points. **Right**: embryo lethality. Surviving embryos were counted at 1 dpf and 4 dpf. Survival was plotted and Log-rank test was done to access differences between groups. Non significant (ns) p > 0.05; * p < 0.05; ** p < 0.01; *** p < 0.001. The number of embryos that were analyzed are indicated in parentheses.

Altogether, these results indicate that co-injection of the OP rescued the developmental defects induced by a pathogenic, basally activated Shp2a variant, while it had no effect on WT embryos.

## Discussion

Our findings provide several insights in the interaction of phosphopeptides with SH2 domains, and in particular, with the N-SH2 domain of SHP2, and on the suitability of this recognition units as therapeutic targets.

Soon after their discovery, the affinities of SH2 domains for their binding partners *(i.e.* the dissociation constants) were considered to fall in the 10-100 nM range [Pawson 1995]. However, it turned out that most of the binding studies performed in that period were affected by experimental artifacts, leading to an overestimation of the binding affinities [Ladbury 1995, Kuriyan 1997]. A reassessment of the affinity values led to a commonly accepted range in the order of 0.1 to 10 μM [Kuriyan 1997, Machida 2005, Wagner 2013, Marasco 2020b]. Such moderate affinities are considered to be crucial for allowing transient association and dissociation events in cell signaling. Consistently, SH2 domains artificially engineered to reach low nanomolar affinities for phosphorylated sequences (known as superbinders) [Kaneko 2012], by increasing the domain affinity for the pY residue, have detrimental consequences for signal transduction. However, micromolar binding affinities make SH2 domains in general less than optimal therapeutic targets. In the case of the N-SH2 domain of SHP2, literature results on the affinity for the IRS-1 pY 1172 peptide were contradictory, with dissociation constants varying by three orders of magnitude [Case 1994, Sugimoto 1994, Keilhack 2005]. Here, we showed that, at least for this peptide, the dissociation constant is in the low nM range. For the N-SH2 domain, similar affinities have been reported also for the GRB2-associated-binding protein 1 (Gab1), pY 627 [Koncz 2001], and Gab2, pY 614 [Bonetti 2018]; several other peptides have a dissociation constant within an order of magnitude of that of IRS-1 pY 1172 [Anselmi 2020, Marasco 2020a]. In addition, in the present study, we were able to further improve the affinity with respect to the parent peptide. Therefore, the N-SH2 domain of SHP2 might constitute an exception in the panorama of SH2 domains, regarding the binding affinity. In most cases, interaction of phosphopeptides with SH2 domains is dominated by the hydrophobic effect (except for the pY pocket). The N-SH2 domain of SHP2 has a peculiar KxK motif in the region of the BG loop pointing towards the binding groove, which can interact electrostatically with basic residues present in the peptide sequence at positions +2 and +4 [Anselmi 2020]. Therefore, by contrast to the superbinders, the high binding affinity of the N-SH2 domain is a result of additional interactions in the selectivity-determining region, and not in the pY pocket. Indeed, our data showed that the pY phosphate contributed less than 30% of the standard binding free energy. This finding is comparable to what has been reported for other SH2 domains [Waksman 2004].

Our data also showed that residue -2 contributes significantly to the binding affinity. Indeed, while the specificity of most SH2 domains is determined by residues C-terminal to the pY, peptide library and array studies have shown that, contrary to most other SH2 domains, the N-SH2 domain of SHP2 has specific preferences for position -2 [Anselmi 2020]. This peculiarity is due to the fact that, in place of the commonly conserved arginine at position 2 in the first α-helix (αA2), the N- SH2 domain of SHP2 has G13. Consequently, a hydrophobic peptide residue at position −2 can insert in the space left free by the missing protein side chain and interact with V14 αA3 and with the phenol ring of pY, stabilizing its orientation and the overall complex [Anselmi 2020].

A preference of the N-SH2 domain of SHP2 for hydrophobic residues at positions +1, +3 and +5 is well established. These side chains insert in the groove on the surface of the domain and interact with exposed hydrophobic patches [Anselmi 2020]. Now our data demonstrate that the bulky, aromatic side chain of tryptophan at position +5 is 10 times better (in terms of dissociation constant) than the leucine residue, which is present in high-affinity natural sequences such as those of IRS- 1, Gab1 and Gab2. Overall, these data confirm that the phosphopeptide sequence in the -2 to +5 stretch contributes significantly to the binding affinity. In principle, highly specific binding should be possible.

In general, SH2 domains are only moderately discriminating for binding target sequences, and a range of residues is tolerated at each site [Kuriyan 1997, Waksman 2004]. Consequently, nonspecific tyrosine phosphorylated sequences are usually bound only 10- to 100-fold more weakly than specific targets [Bradshaw 1998, Waksman 2004, Machida 2005, Wagner 2013]. Indeed, additional specificity is often provided by tandem SH2 domains [Waksman 2004]: two closely- spaced tyrosine phosphorylated motifs bind to tandem SH2 domains with 20- to 50-fold greater affinity and specificity compared with the binding of a single SH2 domain with a single tyrosine phosphorylated motif [Wagner 2013]. SHP2 and its SH2 domains are no exception in this case, as peptide library and array studies, together with the sequences of known natural binding partners, showed a significant variability in the sequences of peptides bound by SHP2 [Anselmi 2020]. However, our results indicate that some peptides (like those developed here) can bind specifically to a single SH2 domain. Among an array of 97 human SH2 domains, we found some interference only with adapter protein APS (also called SHP2B2). The structure of the APS SH2 domain in complex with a cognate protein shows that the phosphorylated sequence binds in a folded, kinked conformation, rather than in the usual extended binding mode [Hu 2003]. This observation should facilitate the further development of our peptides, to avoid the unwanted interaction with APS, without affecting the affinity for the target N-SH2.

Finally, it is worth mentioning that approximately one order of magnitude in affinity was lost by substituting the pY residue with the non-dephosphorylatable mimic F_2_Pmp. This finding is consistent with previous studies showing that F_2_Pmp is tolerated differently by various SH2 domains, and its insertion in place of pY in a phosphopeptide sequence can lead either to a loss or to an increase in affinity, by approximately one order of magnitude [Burke 2006]. Further optimization of this aspect is warranted, but the dissociation constant of our non- dephosphorylatable peptide remained in the nM range.

The non-dephosphorylatable peptide allowed novel experiments on several aspects of SHP2 function and regulation. As discussed in the introduction, in the autoinhibited state of SHP2, the N- SH2 binding groove is closed, apparently making phosphopeptide association impossible. By contrast, the N-SH2 binding site is open in the structure of the isolated N-SH2 domain or of active SHP2. Consequently, it has been hypothesized that mutations destabilizing the closed state and favoring SHP2 activation could lead to an increase binding affinity [Bocchinfuso 2007]. This idea is indirectly supported by the fact that basally activated mutants require lower concentrations of SH2 domain-binding phosphopeptides to reach full activation [Keilhack 2005; Bocchinfuso 2007, Martinelli 2008, LaRochelle 2018]. However, no direct measurements of phosphopeptide binding to different SHP2 variants had been reported until now. Our data directly demonstrate that the affinity for phosphopeptides of activated variants of SHP2 can increase by a factor of 20, reaching the same value as the isolated domain in the most active mutants (Figure 6). This consequence of pathogenic mutations adds to the increase in basal activity and might be the main responsible for hyperactivation of signaling pathways modulated by SHP2. Interestingly, in view of possible therapeutic applications, in a cellular environment, our peptide would act as an effective inhibitor of the PPIs of mutant, hyperactivated SHP2, while it would have a much lower effect on the WT protein. This behavior is the exact opposite of what has been observed for allosteric inhibitors, such as SHP099, which have a significantly impaired activity in pathogenic variants of SHP2 [LaRochelle 2018; Tang 2020].

A second link between SHP2 activity and binding functions is provided by literature data indicating for SHP2 interactors a role both as ligands of the SH2 domains and as substrates of the catalytic PTP domain, often with the same phosphorylated sequence. Examples include IRS-1 [Sugimoto 1994, Noguchi 1994], Gab1 [Cunnick 2001, Zhang 2002], Gab2 [Gu 1997], PDGFR [Rönnstrand 1999, Sugimoto 1992], PD-1 [Marasco 2020a] and SHPS-1 [Fujioka 1996, Takada 1998]. We also showed here that our modified sequence can be dephosphorylated. These data indicate the possible presence of a still uncharacterized feedback mechanism to regulate SHP2 signaling. Using our non- dephosphorylatable peptide, we could demonstrate that a N-SH2-binding sequence associates to the catalytic PTP domain, too, although with a lower affinity. This finding suggests that it might be possible to develop double-edged sword molecules, able to inhibit both the catalytic activity and the PPIs of SHP2.

Phosphorylated sequences cause SHP2 activation by binding to the N-SH2 domain and inducing or stabilizing a domain conformation that is incompatible with the N-SH2/PTP interaction. In principle, it is possible that different phosphopeptide sequences do not have the same aptitude for causing or favoring the conformational transition of the N-SH2 domain needed for SHP2 activation. In this case, the binding affinity and activating potential of phosphopeptides would not be strictly coupled. Some literature data indicate that this might be the case. For instance, the sequences corresponding to pY546, pY895 and pY1222 of IRS-1 (rat ortholog numbering) [Sugimoto 1994, Case 1994], or the artificial sequences AALNpYAQLMFP and AALNpYAQLWYA [Imhof 2006] have similar dissociation constants for the N-SH2 domain (within a factor of 2), but the concentrations of these phosphopeptides needed for full activation of SHP2 differ by orders of magnitude. The interpretation of these studies is complicated by the fact that in principle these sequences could be dephosphorylated by SHP2, to different extents, during the activation experiments. Our data show that a concentration of nonhydrolyzable phosphopeptide that almost completely saturates the N-SH2 domain causes only partial activation. While an inability of this specific sequence to favor activation cannot be ruled out, it is possible that partial activation is caused by inhibition of SHP2 activity due to peptide association to the PTP domain. In any case, even if activation of SHP2 without proper PPIs is inconsequential [Saxton 1997, Shi 1998, Higashi 2002], it is important to note that the molecules developed here can inhibit the association of SHP2 to its partners, without causing complete activation, particularly for the WT protein.

Inhibition of PPIs, particularly using peptides, is currently a hot area of pharmaceutical research. For the RAS/MAPK pathway alone, at least 30 inhibitors of PPI have been developed and several of them are undergoing clinical trials [García-Gómez 2018]. However, no studies on SHP2 have been reported, notwithstanding the central role of this phosphatase in the pathway. Peptides are particularly appealing for the inhibition of PPIs, where large interfaces are involved, which are difficultly targeted selectively by small molecules. An increasing number of drugs based on peptides or peptidomimetics is progressing in the drug development pipeline [Henninot 2018]. Possible challenges in the therapeutic applications of peptide-based molecules are their rapid degradation and a poor cell uptake, particularly for highly charged sequences [Henninot 2018]. Here we successfully overcame the first hurdle, thanks to the introduction of non-natural amino acids. Several studies have demonstrated that efficient intracellular delivery of phosphopeptide mimics is possible, for instance by conjugation to cell-penetrating sequences [Kertesz 2006, Ye 2007, Choi 2009, Nasrolahi Shirazi 2013, Cerulli 2020b]. Optimization of the cell uptake of our molecules, through different strategies, is currently underway.

Our *in vivo* findings on zebrafish embryos are very promising in view of potential applications, particularly considering that the peptide is more effective on activating mutants than on the WT protein, contrary to allosteric inhibitors such as SHP099 [LaRochelle 2018, Tang 2020]. Indeed, besides their possible use as a research tool to study the role of PPIs in the function of SHP2, and regulation of the pathways controlled by this protein, including RAS/MAPK and PI3K/AKT signaling, the reported peptides constitute lead compounds for the development of new drugs against malignancies driven by *PTPN11* mutations, such as JMML, AMoL, and ALL, also considering that allosteric inhibitors have low activity against basally activated SHP2 variants [Larochelle 2018, Tang 2020]. Finally, another possible field of therapeutic application is represented by rare diseases such as NS and NSML, which are caused by activating mutations of *PTPN11* (against which the available allosteric inhibitors are poorly active) and cause several severe postnatal, evolutive clinical manifestations, particularly hypertrophic cardiomyopathy [Tartaglia 2010].

## Materials and methods

### Materials

Fmoc (9-fluorenylmethyloxycarbonyl)-amino acids were obtained from Novabiochem (Merck Biosciences, La Jolla, CA). Rink amide MBHA resin (0.65 mmol/g, 100-200 mesh) was purchased from Novabiochem. All other protected amino acids, reagents and solvents for peptide synthesis were supplied by Sigma–Aldrich (St. Louis, MO). The LB medium components, all the reagents used to prepare the buffers and the Bradford reagent were purchased from Sigma Aldrich. Tris(2- carboxyethyl)phosphine (TCEP) was obtained from Soltec Ventures, Beverly, MA, USA. Spectroscopic grade organic solvents were purchased from Carlo Erba Reagenti (Milano, Italy). Cell culture media growth factors and antibodies were purchased from VWR International PBI (Milan, Italy), EuroClone (Milan, Italy), Promega (Madison, WI, USA), Invitrogen (Carlsbad, CA, USA), Cell Signaling (Danvers, MA, USA), Sigma-Aldrich (Saint Louis, MO, USA), and Santa Cruz Biotechnology (Dallas, TX, USA).

### Peptide synthesis

The solid-phase peptide synthesis of the analogs described in this manuscript was performed on the Rink Amide MBHA resin using standard Fmoc chemistry protocols. using standard Fmoc chemistry protocols. The deprotection of the Fmoc group was performed with a 20 % piperidine solution in N,N-dimethylformamide. The single coupling steps were carried out in the presence of HBTU/HOBt/DIPEA. The N-termini of the different peptides were manually acetylated using a mixture of acetic anhydride (Ac2O) and DIPEA in DMF. For the fluorescent analogs, the introduction of the carboxyfluorescein probe was carried out activating 5(6)-carboxyfluorescein in the presence of HBTU/HOBT/DIPEA, repeating the acylation step twice. The fluorescent analogs were obtained as a mixture of isomers. At the end of the synthesis, each peptide was cleaved from the resin using a mixture of TFA, TIS, and water in 95:2.5:2.5 ratio. The solution was concentrated under a flow of nitrogen, and the crude peptide precipitated by addiction on diethyl ether. The crude peptides were purified by flash chromatography on Isolera Prime chromatographer (Biotage, Uppsale, Sweden) using a SNAP Cartridge KP-C18-HS 12g or preparative RP-HPLC on a Phenomenex C18 column (22.1x250 mm, 10 μm, 300 Å) using an Akta Pure GE Healthcare (Little Chalfont, UK) LC system equipped with an UV-detector (flow rate 15 mL/min) and a binary elution system: A, H_2_O; B, CH_3_CN/H_2_O (9: 1 v/v); gradient 25-55% B in 30 min. The purified fractions were characterized by analytical HPLC-MS on a Phenomenex Kinetex XB-C18 column (4.6 x 100 mm, 3.5 μm, 100 Å) with an Agilent Technologies (Santa Clara, CA) 1260 Infinity II HPLC system and a 6130 quadrupole LC/MS. The purity and the characterization data are reported in Table S2. Peptides were dissolved in DMSO to obtain stock solutions between 1 and 1.5 mM. The exact concentration was obtained by UV measurements, exploiting the signal of carboxyfluorescein for the labeled peptides and of pTyr, Tyr and Trp for the unlabeled peptides. To this end, CF-labeled peptides were diluted from the stocks (1:100) in buffer (pH 9), and their concentration was calculated from the CF signal at 490 nm using a molar extinction coefficient of 78000 M^-1^ cm^-1^ [Esbjörner 2007]. Unlabeled peptides were diluted 1:100 in a pH 7.4 buffer; molar extinction coefficients of Tyr, Phe and Trp were taken from reference [Pace 1995], while molar extinction coefficient of pY was taken from [Bradshaw 2003].

### Protein expression and purification

The human esaHis-tagged *PTPN11* (residues 1-528) cDNA was cloned in a pET-26b vector (Novagen, MA, USA). Nucleotide substitutions associated with NS or leukemia were introduced by site-directed mutagenesis (QuikChange site-directed mutagenesis kit; Stratagene, CA, USA). A construct containing the cDNA encoding the isolated PTP domain preceded by the C-SH2 domain (residues 105-528) was generated by PCR amplification of the full-length wild-type cDNA and subcloned into the pET-26b vector (SHP2_Δ104_). A similar procedure was followed for the constructs of the N-SH2 (residues 2-111), C-SH2 (109-217) and PTP (212-528) domains, and of the N-SH2/C- SH2 tandem (2-217). Primer sequences are available upon request.

Recombinant proteins were expressed as previously described [Martinelli 2012], using *E. coli* (DE3) Rosetta2 competent cells (Novagen). Briefly, following isopropyl β-D-1- thiogalactopyranoside (Roche) induction (2 hr at 30 °C, or overnight at 18 °C), bacteria were centrifuged at 5,000 rpm, 4 °C for 15 minutes, resuspended in a lysozyme-containing lysis buffer (TRIS-HCl 50 mM, pH=8.5, NaCl 0.5 M, imidazole 20 mM, tris(2-carboxyethyl)phosphine (TCEP) 1mM, lysozyme 100 mg/ml, 1 tablet of complete protease inhibition cocktail) and sonicated. The lysate was centrifuged at 16,000 rpm, 4 °C for 30 minutes. The supernatant was collected, and the protein of interest was purified by affinity chromatography on a Ni-NTA column (Qiagen, Hilden, Germany), using a TRIS-HCl 50 mM, NaCl 0.5 M, TCEP 1 mM buffer containing 100 mM or 250 mM imidazole, for washing and elution, respectively. To remove imidazole, the samples were then dialyzed in a 20 mM TRIS-HCl (pH 8.5) buffer, containing 1 mM TCEP and 1 mM EDTA and 50mM NaCl (or 150 mM NaCl if no further purification steps followed). Full length proteins and the SHP2_Δ104_ construct were then further purified by sequential chromatography, using an Äkta FPLC system (Äkta Purifier 900, Amersham Pharmacia Biotech, Little Chanfont, UK). The samples were first eluted within an anion exchange Hi-Trap QP 1ml-column (GE Helathcare, Pittsburgh, PA, USA); the elution was carried out using TRIS-HCl 20 mM (pH 8.5) in a NaCl gradient from 50 to 500 mM. The most concentrated fractions were then eluted in a gel filtration Superose column using TRIS-HCl 20 mM buffer containing NaCl (150 mM) as mobile phase. Sample purity was checked by SDS PAGE with Coomassie Blue staining and resulted to be always above 90%. Proteins were quantitated by both the Bradford assay and the UV absorbance of aromatic residues, calculating extinction coefficients according to [Pace 1995]. In general, the two methods were in agreement, but the values derived from UV absorbance were more precise and are reported in the Figures and Tables. The protein samples were used immediately after purification or stored at -20 °C and used within the following week. In this case, after thawing TCEP 2.5 mM was added, the samples were centrifuged at 13,000 rpm for 20 minutes, and the new concentration was re-evaluated. In the few cases where residual apparent absorbance due to light scattering was present in the UV spectra, it was subtracted according to [Castanho 1997].

### Phosphatase activity assays

Catalytic activity was evaluated *in vitro* using 20 pmol of purified recombinant proteins in a 200 µl reaction buffer supplemented with 20 mM *p*-nitrophenyl phosphate (Sigma) as substrate, either basally or following stimulation with the protein tyrosine phosphatase nonreceptor type substrate 1 (PTPNS1) bisphosphotyrosyl-containing motif (BTAM peptide) (GGGGDIT(pY)ADLNLPKGKKPAPQAAEPNNHTE(pY)ASIQTS) (Primm, Milan, Italy), as previously described [Martinelli 2008]. Proteins were incubated for 15 min (SHP2_Δ104_) or 30 min (SHP2) at 30 °C. Phosphate release was determined by measuring absorbance at 405 nm. DiFMUP (6,8-difluoro-4-methylumbelliferyl phosphate) assay was carried out as previously described [Chen 2016], with minor changes. Briefly, reactions were performed at room temperature in 96-well flat bottom, low flange, non-binding surface, black polystyrene plates (Corning, cat. no. 3991), using a final volume of 100 μl and the following assay buffer: 60 mM HEPES, pH 7.2, 75 mM NaCl, 75 mM KCl, 1 mM EDTA, 0.05% Tween-20, 5 mM DTT. Catalytic activity was checked using 1 nM SHP2 and different concentrations of activating peptides. After 45 min at 25 °C, 200 μM of surrogate substrate DiFMUP (Invitrogen, cat. no. D6567) was added to the mix, and incubated at 25 °C for 30 min. The reaction was stopped by addition of 20 μl of 160 μM bpV(Phen) (Potassium bisperoxo(1,10-phenanthroline) oxovanadate (V) hydrate) (Enzo Life Sciences cat. no. SML0889-25MG). The fluorescence was monitored using a microplate reader (Envision, Perkin- Elmer) using excitation and emission wavelengths of 340 nm and 455 nm, respectively.

The ability of SHP2 to dephosphorylate the phosphopeptides was evaluated through a malachite green phosphatase assay (PTP assay kit 1 Millipore, MA, USA). The BTAM peptide and the following monophosphorylated peptides derived from known SHP2 substrates were used for comparison: DKQVEpYLDLDL (GAB1_Y657_), EEENI<u>pY</u>SVPHD (p190A/RhoGAP_Y1105_), and VDADEpYLIPQQ (EGFR_Y1016_) (Primm). SHP2_Δ104_ (2.4 pmol) was incubated with 100 μM of each phosphopeptide (total volume 25 μl) for different times. The reaction was stopped by adding 100 μl of malachite green solution. After 15 min, absorbance was read at 655 nm, using a microplate reader, and compared with a phosphate standard curve to determine the release of phosphate. Data obtained in the linear region of the curve were normalized on the reaction time (1 min).

### Fluorescence anisotropy binding assay

Anisotropy measurements were carried out using a Horiba Fluoromax 4 spectrofluorimeter. For the binding assays, the requested peptide amount (1nM or 0.1 nM) was diluted in buffer (HEPES 10 mM, NaCl 150 mM, EDTA 1 mM, TCEP 1 mM, fluorescence buffer henceforth) and its anisotropy signal was recorded. The peptide was then titrated with increasing protein amounts, until the anisotropy signal reached a plateau at its maximum value, or up to a protein concentration where protein aggregation and consequent light scattering affected the anisotropy values (usually above 1 μM). The measurements of CF-labeled peptides were carried out using an excitation wavelength of 470 nm and collecting the anisotropy values at an emission wavelength of 520 nm. A 495 nm emission filter was used. For the Cy3-labeled peptides, excitation and emission wavelengths of 520 and 560 nm were used. The lowest peptide concentration needed to have a sufficient fluorescent signal (0.1 nM) was used in the binding experiments. Higher concentrations (1 or 10 nM) were used for peptides with lower affinities, and therefore higher *K_d_* values.

The displacement assays were carried out with the same experimental settings. In this case, the labeled peptide-protein complex was titrated with increasing amounts of the unlabeled peptide, following the decrease in anisotropy. Measurements were carried out at the same CF-peptide concentration used for the corresponding binding experiments. Regarding the protein concentration, a compromise between two requirements is needed [Huang 2003]. On one hand, it is desirable to have a significant fraction of the CF-peptide bound to the protein, to maximize the dynamic range in the anisotropy signal, which decreases during the displacement experiment. On the other hand, the protein concentration should be comparable or lower than the dissociation constant of the unlabeled peptide (*K_i_*), to allow a quantitative and reliable determination of its binding affinity. Since several unlabeled peptides had a higher affinity than their fluorescent counterparts, in the displacement assays we used a protein concentration [*P*]*_T_*∼*K_d_*, or in some cases even ∼*K_d_*/2.

The equations used for data fitting are described in the supplementary information appendix.

### SH2 domain microarray

The microarray experiment was conducted by the Protein Array and Analysis Core at the MD Anderson Cancer Center (University of Texas, USA), as previously described [Roth 2019]. Briefly, a library of SH2 domains [Huang 2008] was expressed as GST fusion in *E. coli* and purified on glutathione-sepharose beads. The domains were spotted onto nitrocellulose-coated glass slides (Oncyte Avid slides, Grace Bio-Labs) using a pin arrayer. Each domain was spotted in duplicate. After incubation with a Cy3-P9W5 solution (0.5, 5.0 nM, or 50 nM), fluorescence signals were detected using a GeneTACTM LSIV scanner (Genomic Solutions).

### In silico studies

#### System preparation

The initial structure of the N-SH2 complexed with phosphopeptide P8 (Table 1) was obtained by amino acid substitutions (and deletions) in the crystallographic structure of the protein complexed with the GAB1 peptide (sequence GDKQVE-pY-LDLDLD) (PDB code 4QSY). The obtained complex was then used as the starting structure for subsequent amino acid substitutions in the bound peptide.

#### System equilibration

MD simulations were performed using the GROMACS 2018.2 simulation package [Abraham 2015] and a variant of AMBER99SB force field with parameters for phosphorylated residues [Homeyer 2006]. Water molecules were described by the TIP3P model. All the simulated systems were inserted in a pre-equilibrated triclinic periodic box (15x7x7 nm^3^), containing about 24000 water molecules and counterions to neutralize system total charge. They were relaxed first by doing a minimization with 5000 steepest descent cycles, by keeping protein positions fixed and allowing water and ions to adjust freely, followed by a heating protocol in which temperature was progressively increased from 100K to 300K. The system was then equilibrated for 100 ps in the NVT ensemble at 300 K, using velocity rescaling with a stochastic term (relaxation time 1 ps) [Bussi 2007] and then for 500 ps at constant pressure (1 atm) using the Parrinello-Rhaman barostat (relaxation time 5 ps). Long-range electrostatic interactions were calculated using the particle mesh Ewald method and the cut-off distance for the non-bonded interaction was set equal to 12.0 Å. The LINCS constraint to all the hydrogen atoms and a 2 fs time-step were used.

#### Preparation of the initial configurations for Umbrella Sampling

For each system, a set of initial configurations was prepared by performing a center-of-mass (COM) pulling simulation. The distance between the peptide and N-SH2 domain COMs was constrained with a harmonic force (K=1000 kJ mol^-1^ nm^-2^). Pulling was performed by gradually increasing the value of the equilibrium distance with a constant-rate of 0.0025 nm/ps. The length of each simulation was about 2.5 ns. During the whole simulation, a positional restraint (1000 kJ mol^-1^ nm^-2^) was applied to all heavy atoms in the N-SH2 domain except for atoms in loops around the binding region (residues 30-45, 52-75, 80-100). For the choice of the optimal unbinding pathway, three different directions were tested, corresponding to: i) the vector from the phosphate to the alpha carbon in pY, in the equilibrated complex; ii) the vector defined by the initial positions of the two COMs; iii) the vector perpendicular to the surface of the cavity flanked by the EF and BG loops, passing through the N-SH2 domain center of mass. Among the three different pathways, the third direction encountered less steric occlusion by the EF and BG loops, and was thus selected for further analyses.

#### Umbrella sampling simulations

A set of starting configurations was extracted from the pull-dynamics trajectory saving the peptide-protein center-of-mass distances every 2 Å in the range from 9 to about 40 Å, thus obtaining about 20 windows along the COM distance. The system in each window was preliminarily equilibrated for 1ns with a strong positional restraint (1000 kJ mol^-1^ nm^-2^) to all carbon alpha atoms except for those in loops flanking the binding region (as in the pull simulation), followed by a production run of 150 ns with the restraints. During this stage, a harmonic potential (K=1000 kJ mol^-1^ nm^-2^) was applied on the distance between the two COMs. Additional sampling windows were added every 1 Å along the distance between the two COMs up to a distance of 15 Å. The resulting asymmetric distribution of sampling windows was used to calculate PMF on the production run trajectories. The Weighted Histogram Analysis Method (WHAM) was used, with default settings (50 bins and tolerance of 10^−6^ kJ mol^−1^), using the gmx wham GROMACS tool. The analysis of the simulation was carried out on the 150 ns production dynamics, during which configurations were stored every 0.1 ns. The statistical uncertainty of the obtained PMF was estimated by bootstrapping analysis [Hub 2010].

### Peptide stability in serum and in DMEM

The peptides were dissolved in DMSO (5mg/mL). In eppendorf tubes, 1 mL of HEPES buffer (25 mM, pH = 7.6) was temperature equilibrated at 37 °C before adding 250 μL of human serum and 20 μL of peptide solution; the reaction was followed for 90 minutes. At fixed intervals, 100 μL of the solution were withdrawn and added to 200 μL of absolute ethanol. These samples were kept on ice for 15 minutes, then centrifuged at 13,000 rpm for 5 minutes; the supernatant solutions were analyzed by HPLC and HPLC-MS with 20-60% B gradient in 20 minutes to follow the reaction. In parallel, samples containing peptide, buffer and ethanol only were analyzed. A degradation resistance test was also conducted in DMEM (Dulbecco’s Modified Eagle Medium). The experimental conditions are similar to those described above; the reaction was followed for 72 hours. The enzymatic degradation resistance tests were followed by HPLC using a 5-50% B gradient in 20 minutes.

### In vivo zebrafish rescue experiments

One cell stage zebrafish embryos were injected with a mixture of 120 ng/µl of mRNA encoding either GFP-2A-Shp2-D61G or GFP-2A-Shp2-wt (as a control), with or without OP, at 0.3 µM, 3 µM and 5 µM concentration. Embryos were selected based on proper GFP expression and imaged at 11 hours post fertilization (hpf) in their lateral position using the Leica M165 FC stereomicroscope. Images were analyzed using ImageJ [Schneider 2012], by measuring the ratio of the major and minor axis from a minimum of 31 embryos. Statistical analysis was performed in GraphPad Prism, using the analysis of variance (ANOVA) complemented by Tukey’s honest significant difference test (Tukey’s HSD). To measure the survival of injected embryos, a minimum of 48 embryos per group were grown up to 4 days post fertilization (dpf) and counted at 1 dpf and 4 dpf. Survival curves were plotted using GraphPad Prism, and the differences between samples were determined using the Log-rank (Mantel-Cox) test.

## Acknowledgment

the authors gratefully acknowledge the Protein Array and Analysis Core at the MD Anderson Cancer Center for performing the SH2 array experiments. This work was supported by AIRC Foundation for Cancer Research in Italy (grants IG19171 and IG24940, to L.S. and IG21614, to M.T.), Italian Ministry of Education, University and Research (MIUR, grant PRIN 20157WW5EH_007, to L.S.), Italian Ministry of Health (Ricerca Corrente 2019 and 2020, to M.T.), European Program on Rare Diseases (NSEuroNet, to M.T. and J.d.H.), Partnership for Advanced Computing in Europe (PRACE, grants 2019204928 and 2017174118, to G. B.), which awarded computational resources at CINECA (Italy), and CINECA (grant HP10BL5G4C, to G. B.). L.P. is a recipient of an AIRC research fellowship.

## Abbreviations Used

ALL: acute lymphoblastic leukemia
AmoL: acute monocytic leukemia
CagA: cytotoxicity-associated immunodominant antigen
JMML: juvenile myelomonocytic leukemia
NS: Noonan syndrome
NSML: Noonan syndrome with multiple lentigines
PTK: protein-tyrosine kinase
PTP: protein tyrosine phosphatase
pY: phosphotyrosine
RTK: receptor tyrosine kinase
SH2: Src homology 2
SHP2: SH2 domain-containing phosphatase 2
shRNA: short hairpin ribonucleic acid
WT: wild type

## Supplementary materials

### Supplementary materials and methods

#### Analysis of binding curves

*K_d_* values were obtained fitting the data with the following equation [Van de Weert 2011], which avoids the ne ed for the commonly used (but often unjustified) approximation of the concentration of unbound protein with the total concentration:

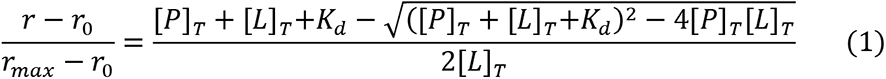

Here, [*P*]_*T*_ and [*L*]_*T*_ are the total protein and ligand concentrations, while *r*, *r_0_* and *r_max_* are the anisotropy values at a given protein concentration, in the absence of protein and when the peptide is completely bound, respectively. When allowed by the experimental conditions (i.e. when [*L*]_*T*_ ≪ *K*_*d*_), this equation was simplified by assuming [*P*]_*T*_ ≅ [*P*] and obtaining:

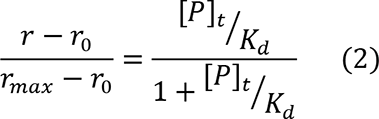

The affinity of unlabeled peptides was determined by competition experiments, in which a sample with fixed total protein and fluorescently labeled peptide concentrations ([*P*]_*T*_ and [*L*]_*T*_) was titrated with the unlabeled peptide, causing displacement of the fluorescent peptide and a decrease in anisotropy. From these data, the *IC_50_* (*i.e.* the total concentration of unlabeled peptide that displaces half of the bound fluorescent analog) was determined, interpolating the displacement curve using a phenomenological Hill equation [Barlow 1989]:

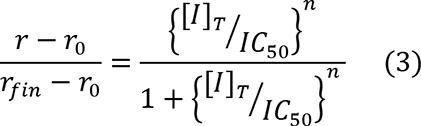

where [*I*]*_T_* is the total concentration of the peptide causing the displacement, and *r_fin_* is the anisotropy corresponding to total displacement, while in this case *r*_0_ is the starting anisotropy, in the absence of displacing peptide.

Successively, the dissociation constant of the unlabeled peptide (*K_i_*) was calculated from the know values of *IC_50_*, *K_d_*, [*P*]*_*T*_* and [*L*]*_*T*_*, as described here below. Our treatment follows that of Nikolovska-Coleska [2004], through a slightly simplified route, and correcting some inaccuracies present in the equations of that article.

In the system where protein (P), ligand (L) and a competitive inhibitor (I) are present, both L and I can form complexes with P (PL and PI, respectively). The following dissociation constants can be defined for the two binding equilibria:

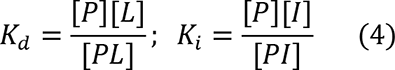

and the following mass conservation laws apply:

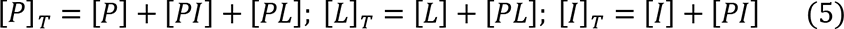

Let’s define [*PL*]_0_ as the complex concentration in the absence of inhibitor. Then, by definition, at the *IC_50_*

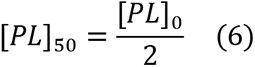

At the *IC_50_*,

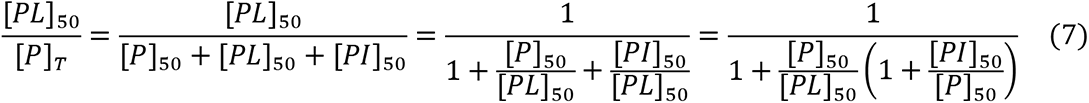

and therefore

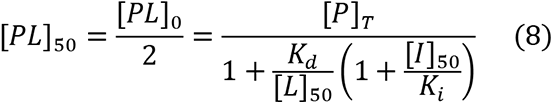

This equation can be inverted to calculate *K_i_*

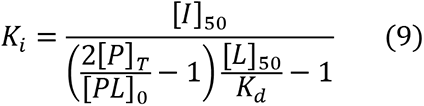

For [*L*]_50_ we can write:

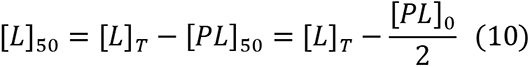

Finally, for [*I*]_50_ we can write:

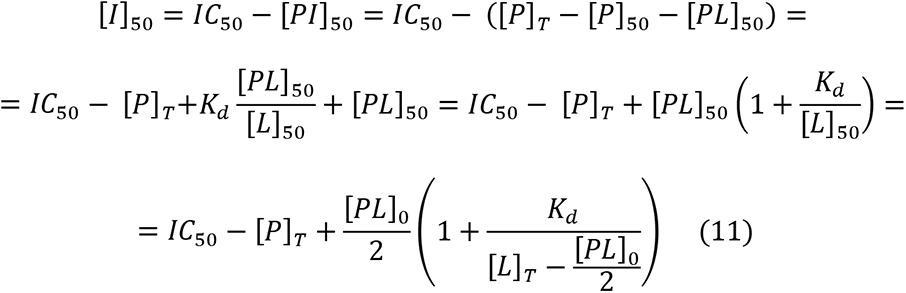

Substituting the above equations in the expression for *K_i_*, we get:

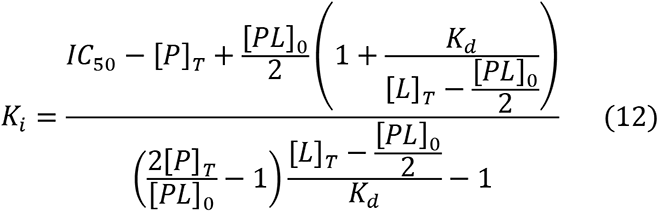

Finally, [*PL*]_0_ can be substituted with the following expression, analogous to Eq. (1):

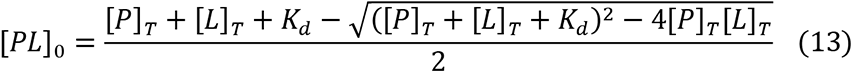

In this way, *K*_*i*_ is expressed as a function of the known quantities *IC_50_*, *K_d_*, [*p*]_T_ and [*L*]_T_, without any approximation.

## Supplementary figures

**Figure S1:**
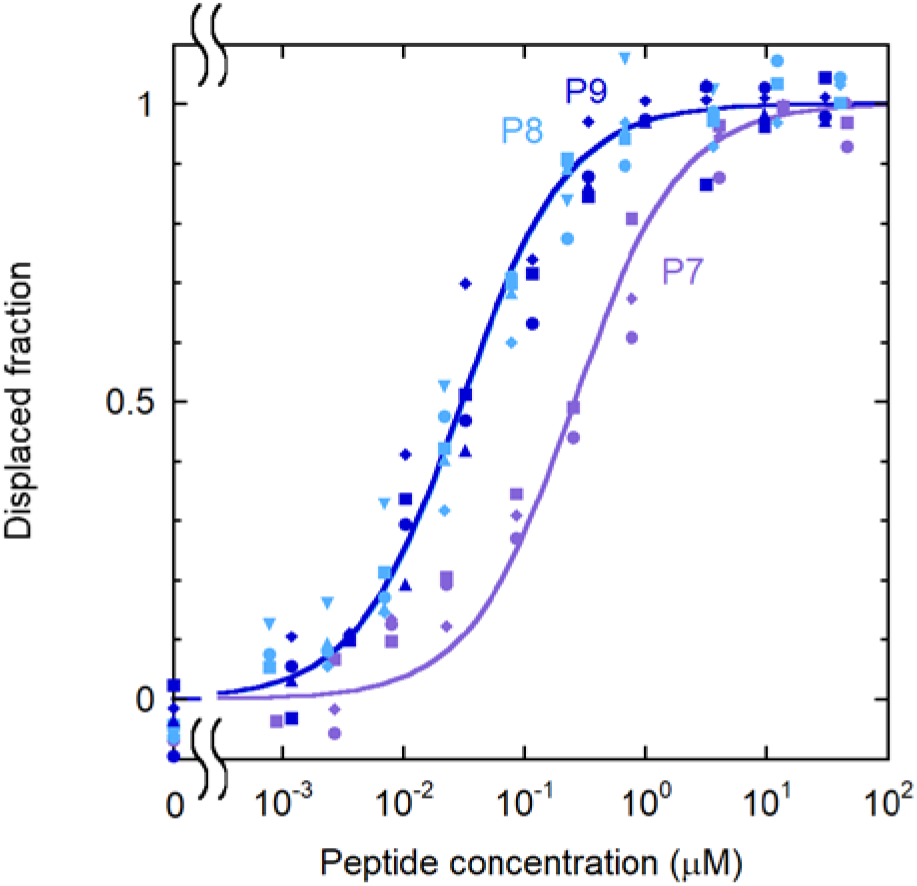
effect of sequence length on the binding affinity. Displacement experiments for analogs of different length. [CF-P9]=1.0 nM; [N-SH2]= 40 nM.

**Figure S2:**
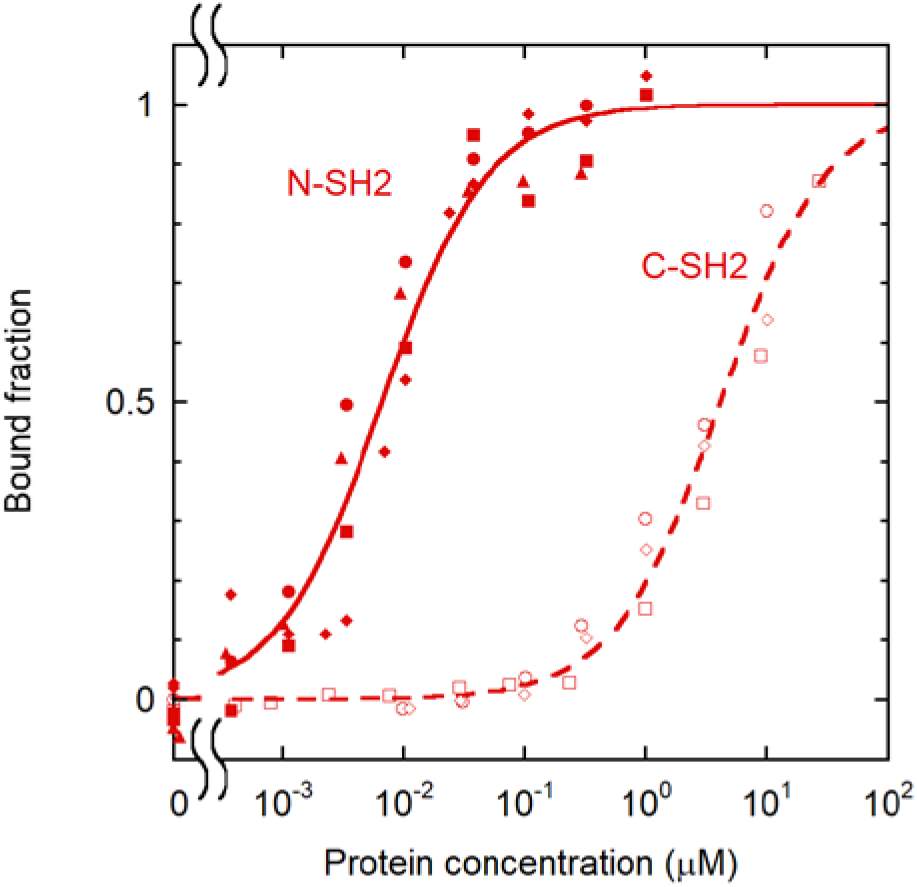
binding selectivity of CF-P9W5 for the two SH2 domains of SHP2. Comparison of the association curves of CF-P9W5 to the N-SH2 and C-SH2 domains of SHP2. Experimental conditions for the N-SH2 binding experiments: see Fig. 5; for the C-SH2 binding experiments: [CF-P9W5]=1.0 nM.

**Figure S3:**
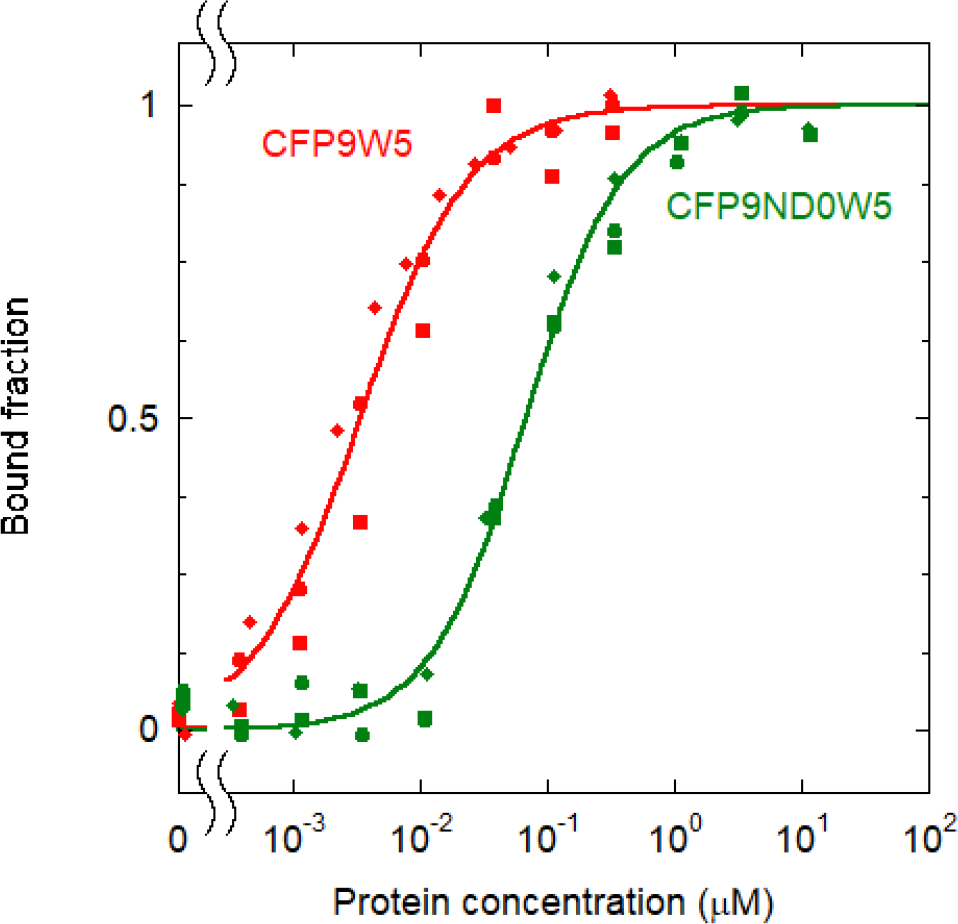
binding of the non-dephosphorylatable peptide CF-P9ND0W5 (or OP) to the N- SH2 domain. For comparison, the curve for CF-P9W5 is also shown. [CF-P9ND0W5]=1.0 nM, [CF- P9W5]=0.10 nM.

**Figure S4.**
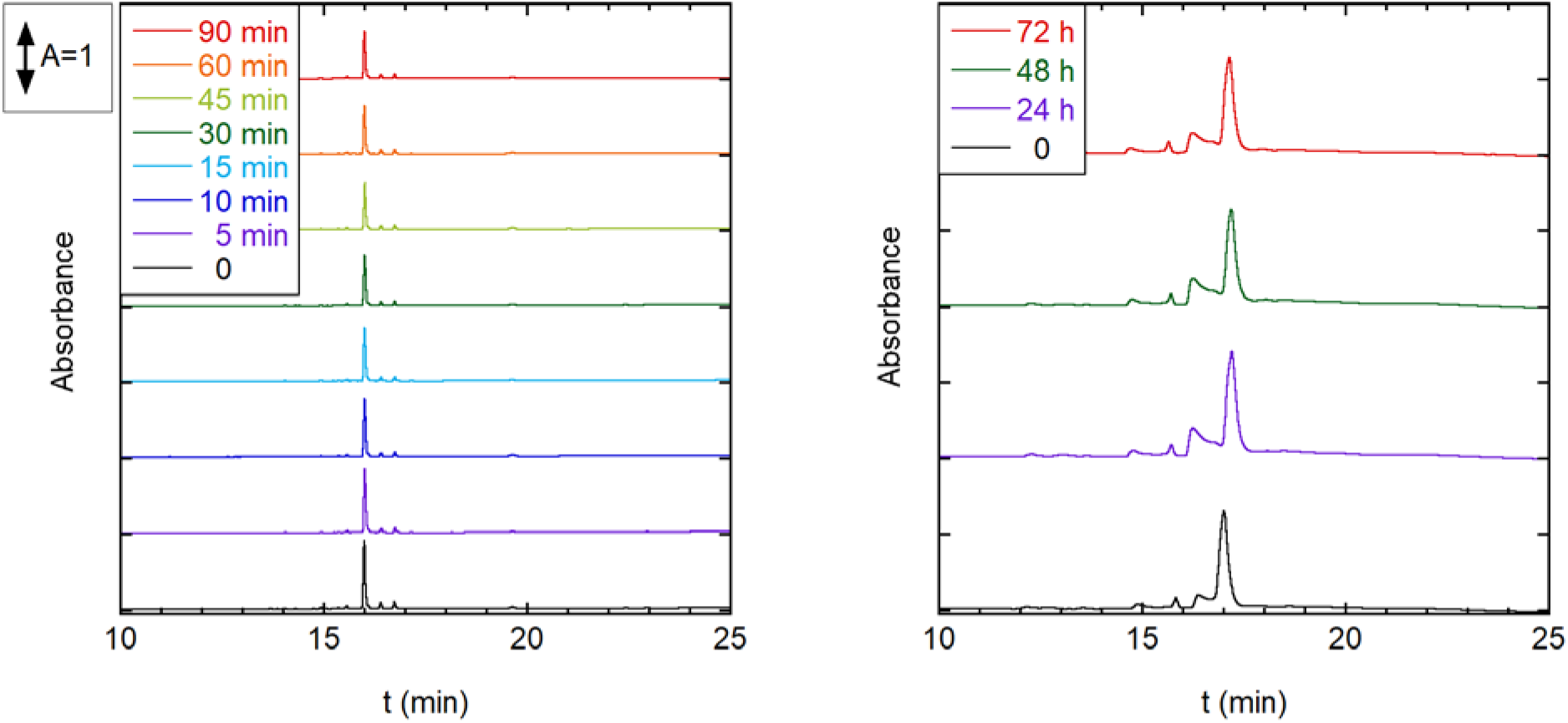
OP resistance to proteolytic degradation in human serum and in DMEM. HPLC profiles of OP (CF-P9ND0W5), after incubation with human serum (left) or DMEM (right), for different times.

**Figure S5:**
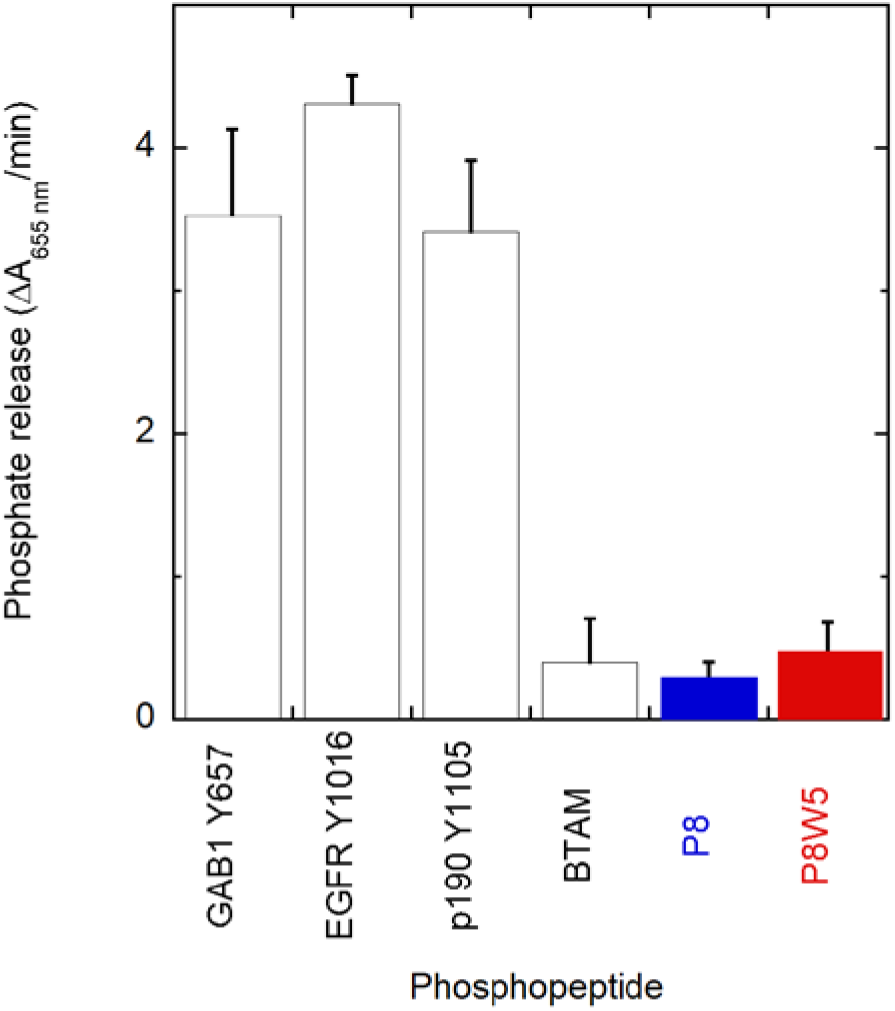
dephosphorylation of P8W5 and other phosphopeptides by SHP2_Δ104_. The following phosphopeptides were used for comparison, in addition to P8 and P8W5. GAB1 Y657 (DKQVEpYLDLDL) p190A/RhoGAP Y1105 (EEENIpYSVPHD) EGFR Y1016 (VDADEpYLIPQQ) BTAM, or bisphosphorylated SHSP-1 TAM1 (GGGGDIT(pY)ADLNLPKGKKPAPQAAEPNNHTE(pY)ASIQTS, with 4 N-terminal G residues) A SHP2 construct lacking the N-SH2 domain (i.e. the first 104 residues, SHP2_Δ104_) was used at a 95 nM concentration. Phosphopeptides were added at a 100 μM concentration and the phosphate released was measured at different times. From the linear region of the phosphate versus time curve, the variation in absorbance at 655 nm in 1 min, due to phosphate release, was calculated and plotted.

**Figure S6.**
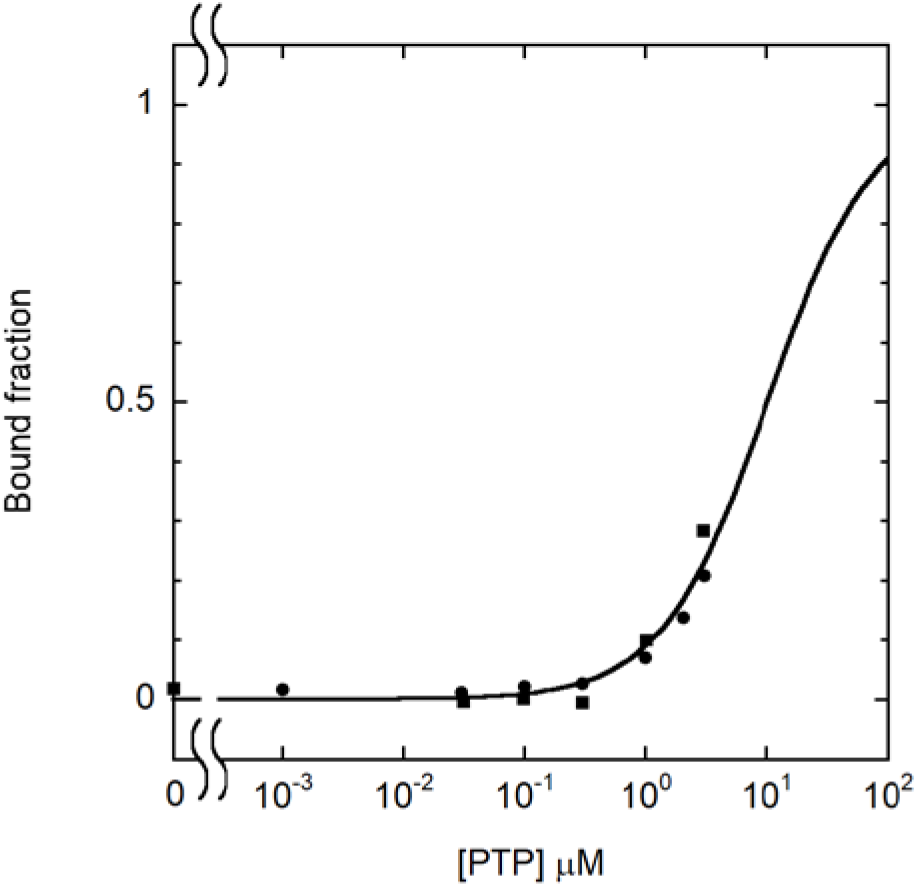
CF-OP association to the PTP domain. [CF-P9ND0W5]=1.0 nM

**Figure S7:**
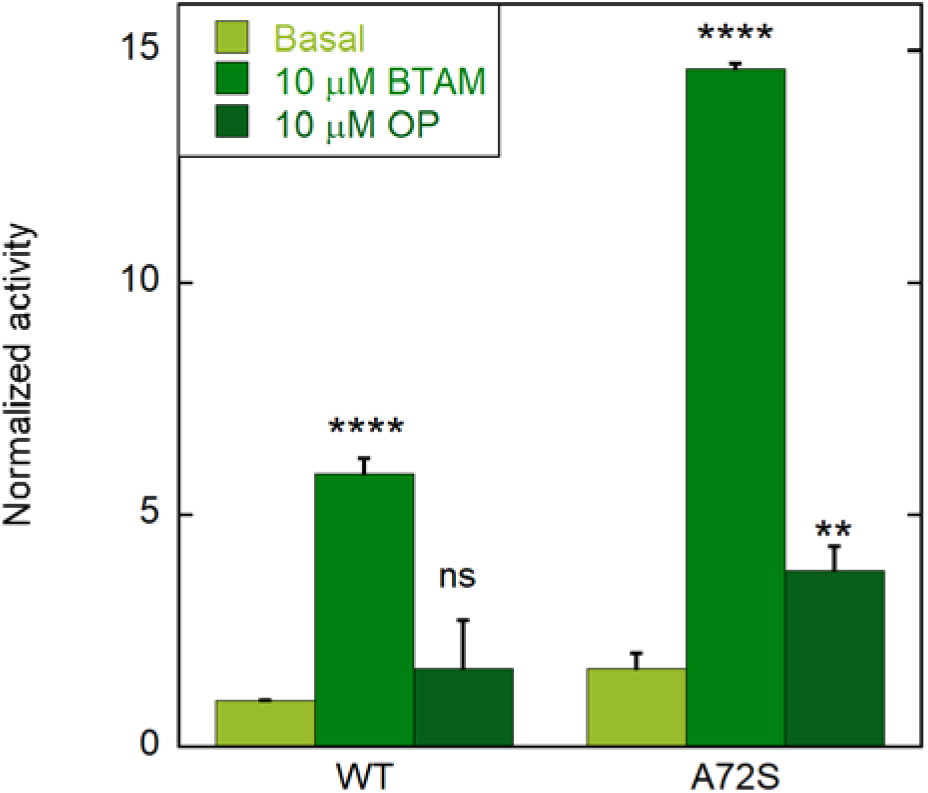
SHP2 activation by the OP. Basal activity is reported in light green, while activities in the presence of 10 µM BTAM or 10 µM OP are shown in green and dark green, respectively. Error bars represent standard deviations. Statistical significance of the difference between the basal activities and the activities in the presence of the peptides, was calculated by ANOVA, complemented by Tukey’s test, and is indicated by asterisks (Non significant (ns) p > 0.05; * p < 0.05; ** p < 0.01; *** p < 0.001; **** p<0.0001)

**Figure S8.**
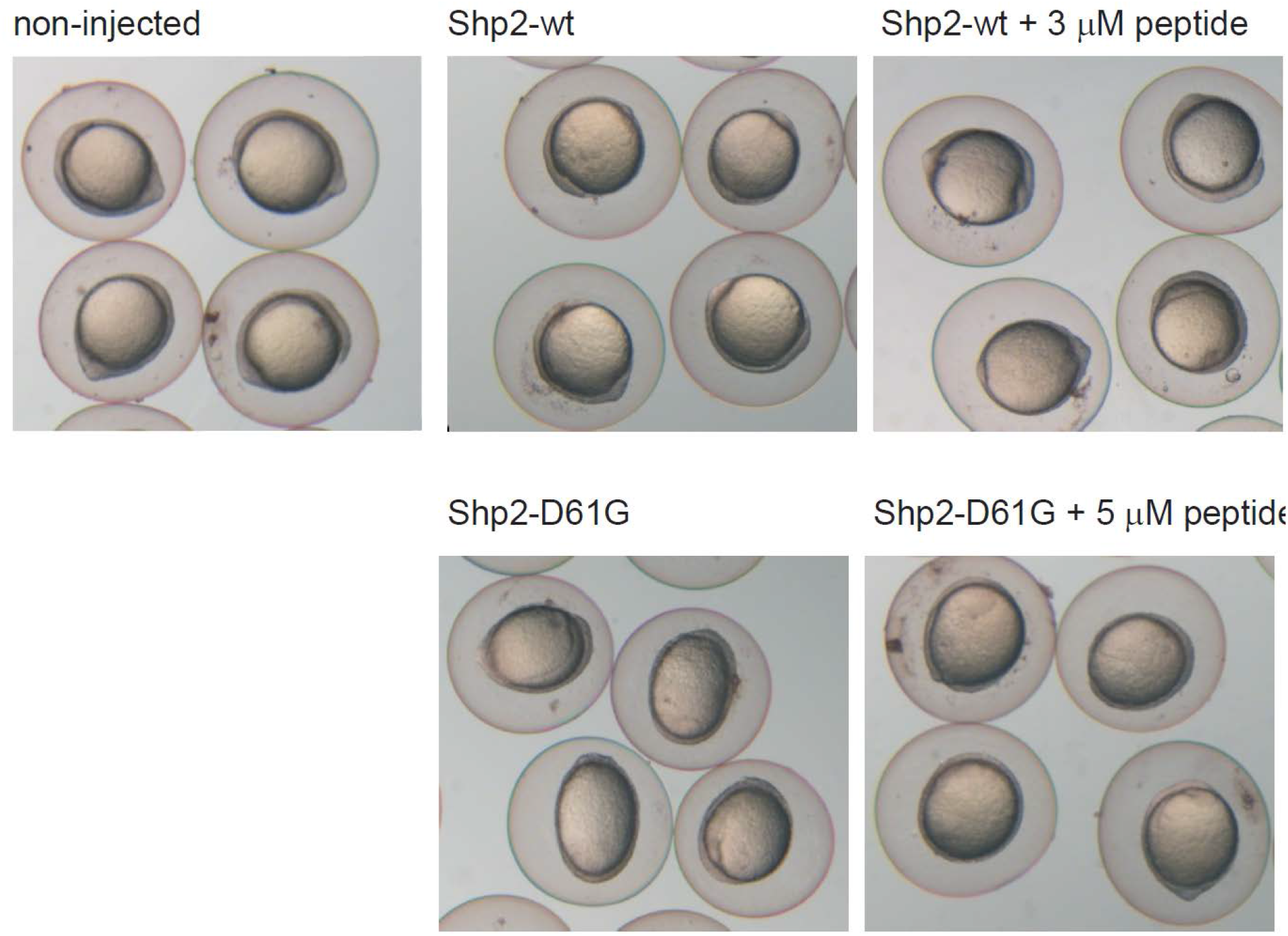
Representative images of zebrafish embryos at 11 hpf. Embryos were injected at the one-cell stage with mRNA encoding GFP-2A-Shp2-D61G or GFP- Shp2-wt with or without peptide at 0.3 µM, 3 µM and 5 µM concentration. Non-injected embryos (ni) were evaluated as a control.

## Supplementary tables

**Table S1.** Literature values for IRS-1 pY1172/N-SH2 domain dissociation constant.

| Reference | Method | K<sub>D</sub> |
| --- | --- | --- |
| Case 1994 | Radioactively labeled peptide | ~1-10 μM |
| Sugimoto 1994 | Surface plasmon resonance | 14±8 nM |
| Keilhack 2005 | Isothermal titration calorimetry | 51 nM |

**Table S2.**
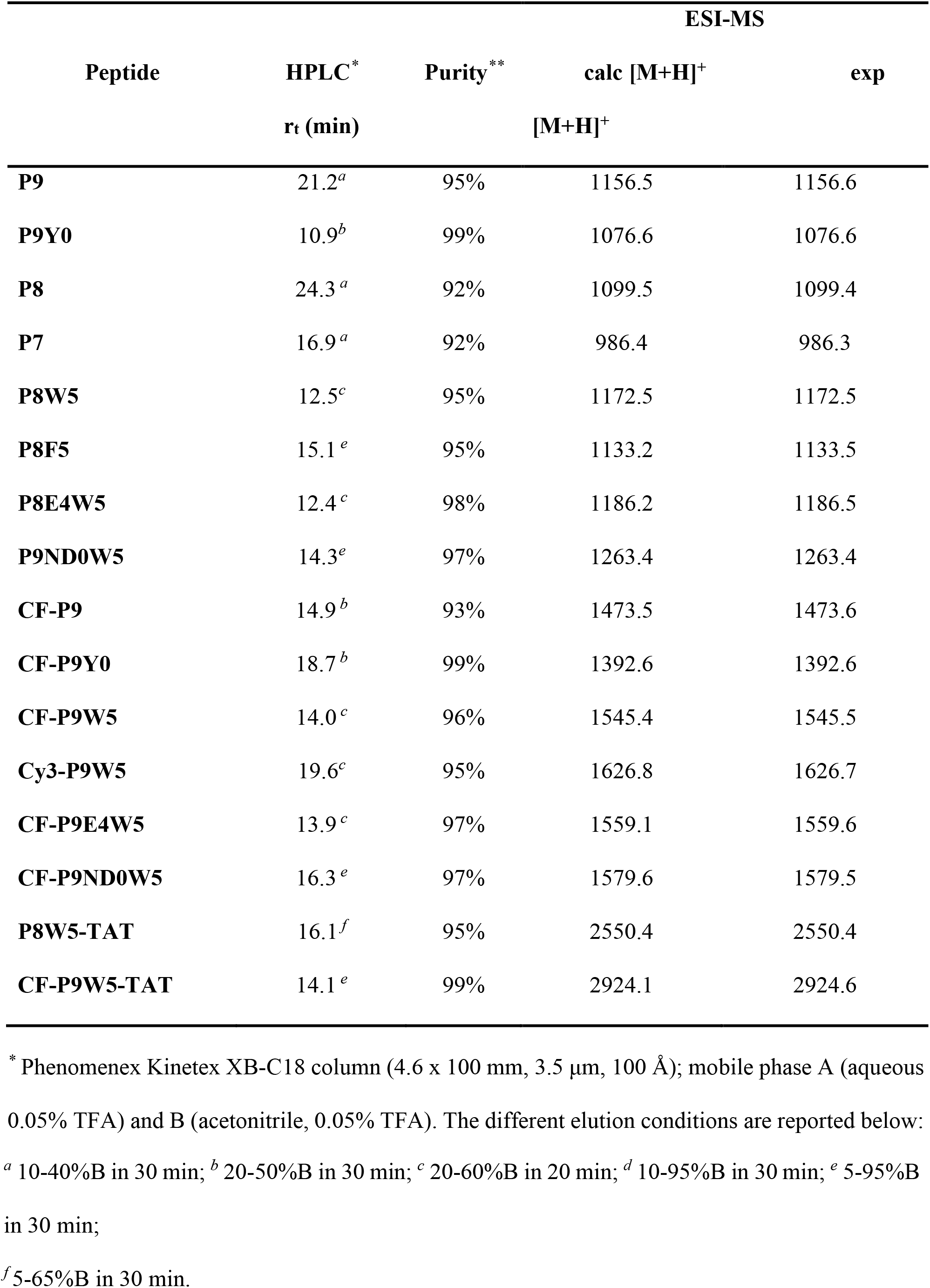
Characterization of the synthesized peptides.

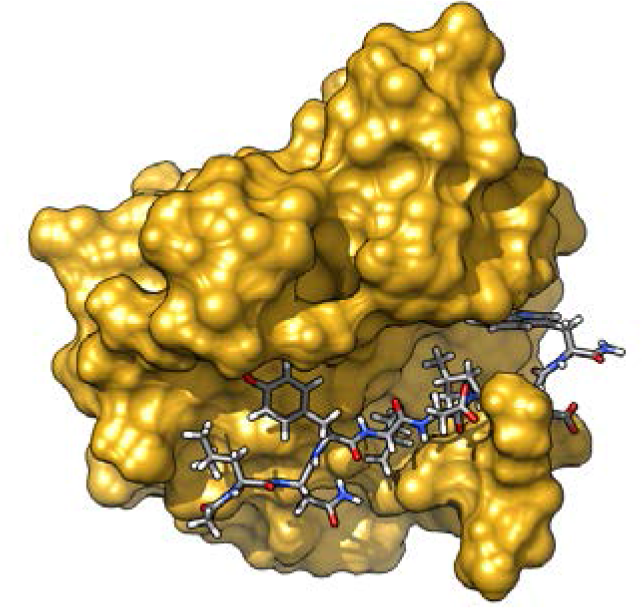
TABLE OF CONTENTS GRAPHICS.

